# Multiomics Interrogation into HBV (Hepatitis B Virus)-Host Interaction Reveals Novel Coding potential in Human Genome, and Identifies Canonical and Non-canonical Proteins as Host Restriction Factors against HBV

**DOI:** 10.1101/2021.03.19.436126

**Authors:** Shilin Yuan, Guanghong Liao, Menghuan Zhang, Yuanfei Zhu, Weidi Xiao, Kun Wang, Caiwei Jia, Qiang Deng, Jian Zhang, Ping Xu, Ronggui Hu

**Author notes:** These authors contributed to this work equally (Shilin Yuan, Guanghong Liao, Menghuan Zhang, Yuanfei Zhu, Weidi Xiao). These authors contributed to this work equally (Qiang Deng, Jian Zhang, Ping Xu, Ronggui Hu).

## Abstract

Hepatitis B Virus constitutes a major threat to global public health. Current understanding of HBV-host interaction is yet limited. Here, ribosome profiling, quantitative mass spectrometry and RNA-sequencing are conducted on a recently established HBV replication system. We have identified multiomic DEGs (differentially expressed genes) that HBV orchestrated to remodel host proteostasis networks. Our multiomics interrogation revealed that HBV induced significant changes in both transcription and translation of 35 canonical genes including PPP1R15A, PGAM5 and SIRT6, as well as the expression of at least 15 non-canonical ORFs including ncPON2 and ncGRWD1, thus revealing an extra coding potential of human genome. Overexpression of these five genes but not the enzymatically deficient SIRT6 mutants suppressed HBV replication while knockdown SIRT6 had opposite effect. Furthermore, the expression of *SIRT6* was down-regulated in patients, cells or animal models of HBV infection. Mechanistic study further indicated that SIRT6 directly binds to mini-chromosome and deacetylates histone H3 lysine 9 (H3K9ac) and histone H3 lysine 56 (H3K56ac), and chemical activation of endogenous SIRT6 with MDL800 suppressed HBV infection *in vitro* and *in vivo*. By generating the first multiomics landscape of host-HBV interaction, our work is thus opening a new avenue to facilitate therapeutic development against HBV infection.

## Introduction

Hepatitis B virus currently infects over 257 million humans worldwide, and chronic HBV infection is the most prominent risk factor for hepatocellular carcinoma (HCC), causing more than 887,000 deaths per year ^1,2^. HBV infection thus constitutes a major global public health threat with yet no complete curing treatment. The compact HBV genome encodes virus DNA polymerase, an X protein (HBx) and virus antigens (HBcAg, HBeAg and HBsAg, respectively). Covalently closed circular DNA (cccDNA) in the form of virus mini-chromosome is central in HBV life-cycle, as it not only shelters the virus from the attack by host pattern recognition factors but also serves as transcriptional template for viral gene expression ^3–6^. Targeting HBV cccDNA reservoir and persistently silencing cccDNA-based transcription are considered essential strategies that should be prioritized to develop HBV curing treatment^7,8^. Current therapies for chronic HBV infection are restricted to type I interferon treatment or nucleos(t)ide analogues (NAs), which target the viral reverse transcriptase. However, interferon therapy has strong side effect, and its efficiency is also limited^9^; while NAs are better tolerated and potent against viremia, it cannot lead to functional cure, that is, the clearance of HBsAg ^9^. Currently, Studies of HBV-host interaction from the perspective of multi-omics is lacking ^10–13^. A comprehensive delineation of the host networks impacted by HBV would advance our current understanding of HBV infection. State-of-art-omics approaches are thus called upon to systematically elucidate the molecular details in HBV-host interaction ^1^.

Ribosome profiling (RiboSeq) is a technique that could determine the sub-population of mRNAs that are actively translated ^14^. Application of RiboSeq has enabled mapping of ribosome footprints on RNAs at nucleotide resolution^15^. The use of harringtonine to arrest ribosomes at the translation initiation site has made it possible to discover novel translational events, including non-canonical open reading frames (ncORFs)^16^. Through the combination of RiboSeq and quantitative proteomic technology such as SILAC (Stable Isotope Labeling by/with Amino acids in Cell culture)^17^, one could digitally assess the abundance of individual proteins under different conditions, and confirm the existence of the translational products of ncORFs. It is interesting to ask how HBV would impact on translation of these ncORFs and whether and what effects the changed translation of the ncORFs might have on HBV replication or virus gene expression.

Sirtuins family proteins (SIRTs) possess the activity of either mono-ADP-ribosyltransferase or deacylases including deacetylase, desuccinylase, demalonylase, demyristoylase and depalmitoylase ^18^. With SIRT6 as a prominent example, the Sirtuin family members have distinct subcellular localization and are known to regulate aging, mitosis, transcription, apoptosis, inflammation, stress responses and metabolism^19^. Previously, SIRTs have been found to associate with HBV replication with sometimes contradicting reports: Deng and colleagues showed all SIRTs activated HBV gene expression ^20^, while Ren et al reported overexpression of SIRT3 mediated transcriptional suppression of the virus genes through epigenetic regulation of HBV cccDNA^21^. One of the remaining questions with the latter study was whether mitochondria-localized SIRT3 might suppress HBV gene transcription only when introduced exogenously. It also remains unclear whether activation of endogenous SIRTs could have any effect on HBV DNA replication or virus gene transcription.

In this work, we profiled HBV-induced changes in ncORFs of host cells, using RiboSeq and SILAC, as well as differentially expressed genes (DEGs) using conventional RNA-sequencing. We found that HBV DNA replication and/or virus gene expression could be significantly altered when PGAM5, PPP1R15A or SIRT6 was ectopically expressed or the translational product of ncPON2 was introduced as well as ncGRWD1. Particularly, we found SIRT6, which has transcription corepressor activity concerning gene silencing, was transcriptionally and translationally down-regulated by HBV, and MDL800, a small molecular agonist of SIRT6^22^, was shown to potently suppress HBV gene expression in both well-established cell^23,24^and mouse models ^25^ for HBV infection.

## Results

### The experimental systems for HBV DNA replication and gene expression

To achieve robust viral replication, two cell-based HBV replication systems were employed: 1) Huh7.5.1 or HepG2 cells co-transfected with pCMV-Cre and prcccDNA to generate HBV cccDNAs in the cells (HBV^+^) (**Fig. S1A**), while the control cells (HBV^−^) were transfected with pCMV-Cre and pCDNA3.1 vector ^23^; 2) HepG2-derived HepAD38 cells that harbors tetracycline-controlled HBV transgene by chromosomal integration ^26^. Expression of viral antigens was then examined at indicated time points (**Fig. S1B-D**). Specifically, the expression of HBsAg and HBeAg were found to reach plateau approximately at 72 h.p.t in the transient transfection system (**Fig. S1B-S1C**) and after 6 days in the Tet-inducible HBV expression system (**Fig. S1D**). Samples were thus collected at these time points, and subjected to RNA-seq, RiboSeq and other analyses.

### Ribosome profiling reveals the translation of 20,533 non-canonical ORFs (ncORFs), some of which was markedly altered by HBV

RiboSeq (ribosome profiling) was performed with Huh7.5.1 cells that were either HBV^+^ or HBV^−^, and ribosome footprints were mapped to the mRNAs of 6,030 genes (**Supplementary file S1**). On average, ribosomes were found to occupy mRNA fragments with lengths peaking at 30-nucleotide width as typically reported before (**Fig. S2A**). Further quality control assays indicated that these data were highly reproducible among all three biological replicates in RiboSeq (**Fig. S2B**).

With the development of ribosome profiling, many non-canonical open reading frames (ncORFs) have been discovered. Translation of these ncORFs often produces functional or non-functional micro-peptides^27,28^. To identify novel translational events in the presence or absence of HBV, an analysis pipeline combined with RiboCode^29^ was built to genome-widely annotate translated ORFs computationally, which defined the ORFs most likely being actively translated based on our RiboSeq data (**Fig. 1A**).

**Figure 1:**
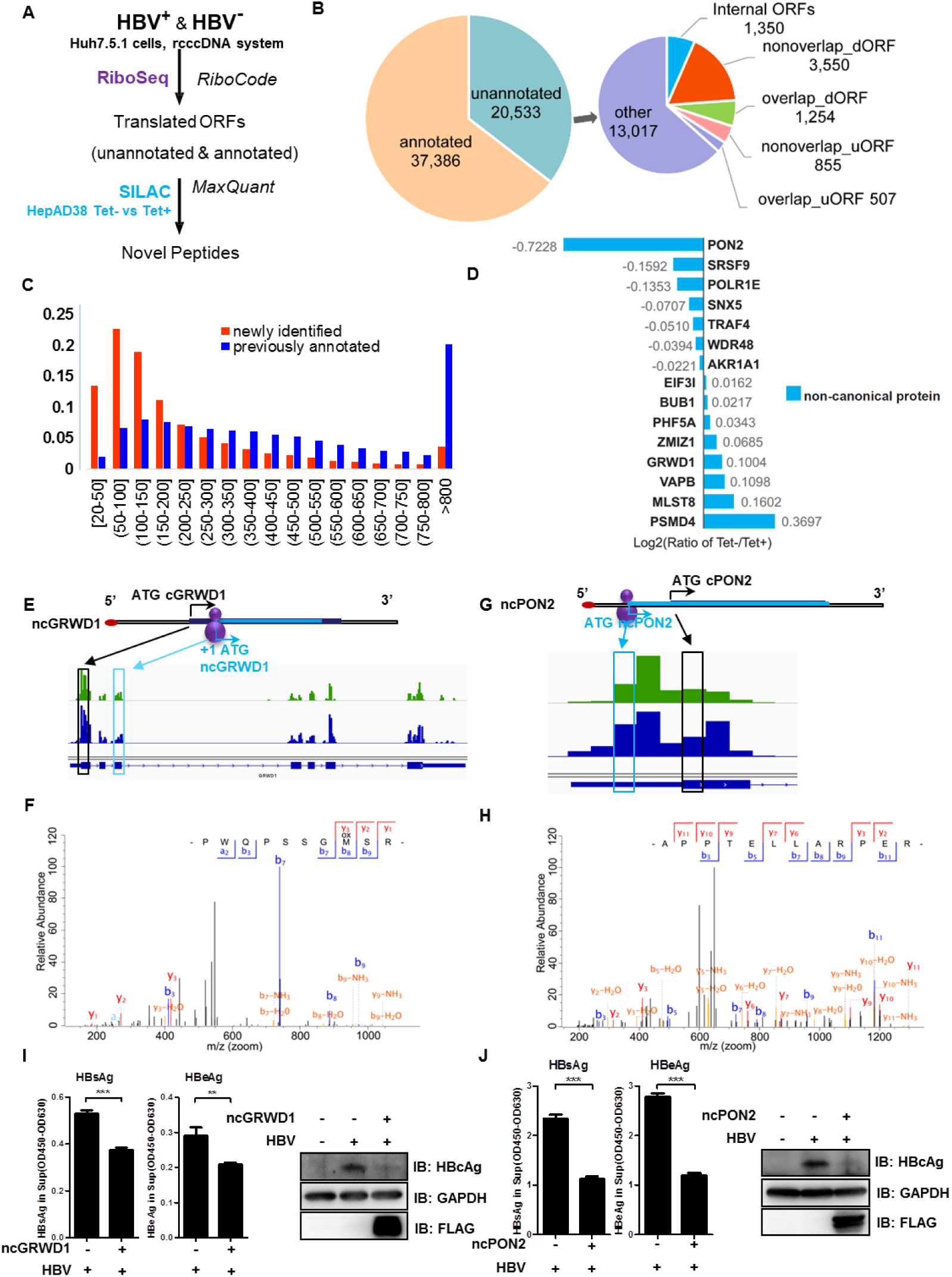
Translation of non-canonical open reading frames (ncORFs) upon HBV replication. (**A**) The workflow for identification of novel ncORFs. (**B**) The subtypes of all translated ORFs identified in this study, see materials and methods for detail. (**C**) The lengths distribution of newly identified ncORFs (blue) and previously annotated ORFs (red). See supplementary materials for detail. (**D**) A list of genes near which the expression of non-canonical ORFs (ncORFs) were altered by HBV. The peptides derived from the translation of these ncORFs were assessed by SILAC, with bar chart to indicate their relative ratios in the presence versus absence of HBV. ^1^ Ribosome occupancy profiles of host ncORFs related to *GRWD1* (E) or *PON2* (G). HBV+ or HBV− of the translational pattern was depicted in green or blue, respectively. The annotated MS/MS spectra of two representative peptides uniquely matched to the translational products of ncGRWD1 (F) or ncPON2 (H) was shown, respectively. The canonical ATG was depicted as black; the non-canonical ATG and the ncORF derived peptide were labeled as cyan. (**I-J**) The inhibitory effect of ncGRWD1 (I) or ncPON2 (J) on expression of HBV antigens was shown. **, p<0.01, ***, p<0.001. ELISA data are presented as bar chart (n = 3).

A total of 57,919 translated ORFs (including ORFs of isoform) were identified using RiboCode in our RiboSeq datasets in HBV^+^ and HBV^−^ groups (**Supplementary file S2**). Among them, 37,386 were perfectly matched to annotated ORFs^29^. The remaining newly identified 20,533 ORFs could be further divided into 6 types (**Fig. 1B**, see materials and methods for more details). These ncORFs were found to encode peptides or proteins of overall lengths markedly shorter than those of the previously annotated canonical ORFs, with over 30% (6,466 out of 20,533) of all translational products less than 100 amino acids in length (**Fig. 1C**).

To further assess the effect of HBV on the translation of host mRNA, we also adopted SILAC^30^ and Maxquant^31^ to detect and quantitate the relative abundances of the products translated from the ncORFs identified above (**Fig. 1A, Fig. S3A**) in HepAD38 cells that had HBV genome integrated in their chromosomes and started to express HBV genes upon the tetracycline withdrawal. After excluding translational products from the in-frame ncORFs that were practically indistinguishable from those of the canonical ORFs, 47 novel peptides were identified and quantified with MaxQuant^31^ against customized sequence database based on RiboCode, with the abundances of many peptides seemed to be altered in host cells upon HBV loading (**Supplementary file S3**). Among them, we sorted 15 novel peptides by the most confident mass spectra result **(Fig.S3B-N)**, and their expression seemed to be significantly perturbed by HBV (**Fig.1D**). There seemed to exist a positive correlation in relative abundance between the translational products of some of the ncORFs and their cognate canonical ORFs, e.g. *PON2, WDR48* and *VAPB* (**Fig.S4A, Supplementary file S10**).

### ncPON2 and ncGRWD1 suppress HBV gene expression

both ncORFs related to *GRWD1* and *PON2* genes and their corresponding translational products were successfully detected in RiboSeq and SILAC analyses, respectively (**Fig. 1E-H**). Replication and expression of HBV in hepAD38 cells were found to increase the expression of a novel translational product started by an internal ATG with +1 frame-shift from canonical *GRWD1* ORF, which we termed as ncGRWD1**(Fig.1E-F)**. GRWD1(Glutamate-rich WD repeat-containing protein 1) was recently identified as a histone binding protein ^32^. Through interaction with DDB1, GRWD1 can be recruited to Cul4B E3 ubiquitin ligase ^33,34^. Interestingly, ectopic expression of ncGRWD1 could suppress the expression of HBc, HBsAg and HBeAg in HBV^+^ cells (**Fig. 1I**).

Notably, among all the novel peptides, a peptide related to the ORF of *PON2* was most significantly down-regulated, and the corresponding translational products was successfully detected by both RiboSeq and SILAC in all three biological replicates, respectively (**Fig. 1D, Fig. S4B**). With HepAD38 (Tet^+^) as the control group that did not express HBV genes except HBs, it was found that the stable replication and expression of HBV genes in HepAD38 cells upon tetracycline withdrawal (Tet^−^) resulted in approximately 40% decrease in the expression of a ncORF in *PON2* gene. This ORF seemed to use an upstream ATG, thus conferring an N-terminal extension to the canonical PON2 protein, which we termed as ncPON2 (**Fig. 1G-H, Fig. S4B**). *PON2* belongs to the paraoxonase gene family, which may act as an antioxidant in cells ^35^. Interestingly, ectopic expression of *ncPON2* suppressed the expression of HBcAg, HBsAg and HBeAg in HBV^+^ cells (**Fig. 1J**).

To take a glimpse into the mechanisms of these two HBV suppressive ncORFs, co-immunoprecipitation coupled with mass spectrometry analysis was performed with ncGRWD1-FLAG and ncPON2-FLAG, with pCDNA3.0-FLAG as a negative control. And GO and pathway enrichment analysis was performed using a web server Metascape (http://metascape.org/) with the identified deemed interactors of ncGRWD1 or ncPON2 (**Supplementary file S4**). The results show that ncGRWD1 could bind to the host machinery participating in metabolism of RNA, translation, RNA splicing and regulation of gene silencing (**Fig. S4C**); while the host proteins interact with ncPON2 was involved in metabolism of RNA, rRNA processing in the nucleus and cytosol, metabolism of lipids and HIV infection (**Fig. S4D**).

Taken together, HBV appeared to alter the translation of some ncORFs including *ncGRWD1* and *ncPON2*, whose translational product could suppress viral gene expression probably through affecting multiple pathways including metabolism of RNA.

### HBV globally impacts on the transcriptional and translational landscapes in host cells

To globally profile the differentially expressed genes (DEGs) in response to HBV, RNA sequencing and ribosome profiling were performed with HBV^+^ Huh7.5.1 cells or the HBV^−^ control cells in parallel (**Fig.2A**). RNA-seq analysis detected the transcripts from 12,547 genes (**Supplementary file S5**), ribosome footprints were mapped to mRNAs of 6,030 genes (**Supplementary file S1**). Across deep-sequencing replicates, our in vivo RNA-seq **(Fig.S5A)** and ribosomal profiling was highly reproducible (**Fig. S2B**).

**Figure 2:**
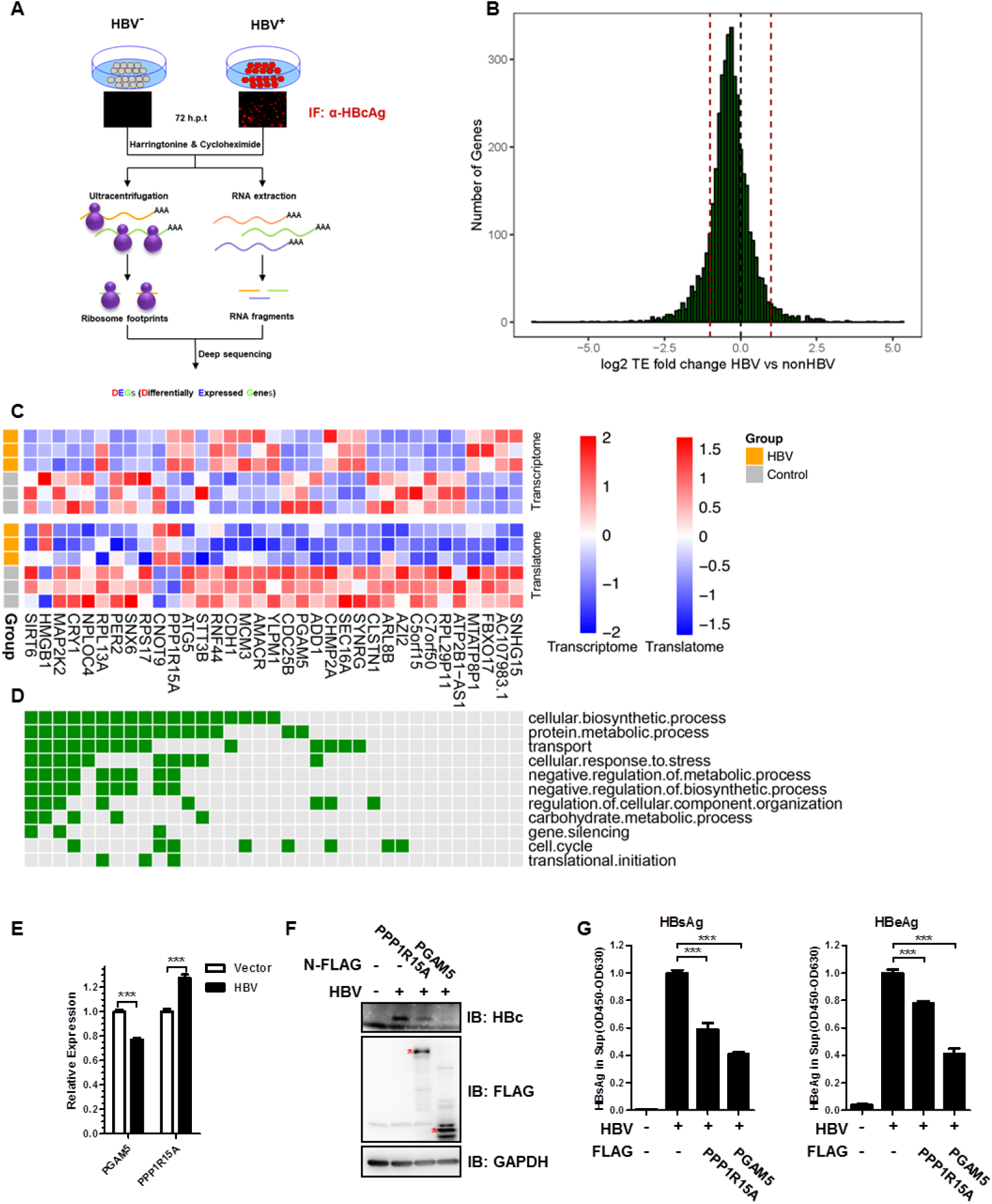
HBV induces significant changes in 35 DEGs at both transcriptional and translational levels (ttDEGs) with PGAM5 and SIRT6 emerging as potential host virus-restricting factors. (**A**) Experimental approaches for transcriptomic and translatomic profiling of host genes upon transfection with the Cre-based rcccDNA system of HBV. (**B**) Differential translational efficiency (TE=RPKM_ribosome profiling_/RPKM_RNA-seq_) in HBV replicating cells versus control cells. (**C**) Heatmap and (**D**) GO annotation of the 35 ttDEGs, whose expression was significantly altered by HBV in both transcription and translation. (**E**) RT-qPCR reveals that HBV down-regulated the transcriptional level of endogenous PGAM5 and up-regulated PPP1R15A transcription. (**F-G**) Reinstatement of the static levels of two representative ttDEGs affected the expression of cellular HBc (F) and secreted HBs and HBe (G). The major bands were marked with asterisks to indicate the theoretical molecular weights of the ectopically expressed ttDEGs. *, p<0.05, **, p<0.01, ***, p<0.001. ELISA data are presented as bar chart (n = 3); qPCR results are presented as bar chart (n = 3).

In general, 324 and 39 genes were significantly up regulated; and 226 and 939 genes were significantly down regulated by HBV in RNA-seq and ribosome profiling, respectively (**Fig.S5B, Supplementary file S6 and S7**). KEGG pathway enrichment analysis identified over-representation of genes for protein processing in endoplasmic reticulum, spliceosome, bacterial invasion of epithelial cells, cysteine and methionine metabolism in the transcriptome-related differential expressed genes (DEGs) (**Figure S5C**); Meanwhile, over-representation of TGF-beta signaling pathway, cellular senescence, adherens junction, N-Glycan biosynthesis, Hippo signaling pathway was identified in the translatome-related DEGs (**Figure S5D**).

To further explore whether HBV impacted on host mRNA translation, we calculated translational efficiency by determining the reads per kilobase of transcript per million mapped reads (RPKM) of coding sequences (CDS) in ribosome profiling versus the RPKM in exons of RNA-seq (RPKM_ribosome profiling_/RPKM_RNA-seq_) in HBV and non-HBV states. The overall translational efficiency of host mRNAs in HBV-transfected cells was lower than in control (**Fig. 2B, Supplementary file S8**), indicating that HBV caused a major shutoff in translation of host genes, a phenomenon also commonly observed with many other viruses ^36,37^.

### HBV induces significant changes in both transcription and translation of 35 host genes including *PPP1R15A* and *PGAM5*

HBV significantly modulated both the transcription and translation of 35 differentially expressed genes (DEGs) (**Fig.2C**). Gene ontology analysis indicated that over half of these 35 DEGs were involved in either biosynthetic or metabolic processes (**Fig.2D**). Among them, many genes such as *SIRT6*, *HMGB1*, *MAP2K2*, *CRY1* and *PER2* were key regulatory factors or functional components in multiple biological processes, including cellular biosynthetic process, protein metabolic process, negative regulation of metabolic process, circadian clock, and negative regulation of biosynthetic process and transport. NPLOC4, RPS17 and PPP1R15A, however, were of known roles in regulating translation (**Fig.2D**). These DEGs were thus serving as the crucially important nodes of multiple metabolic and signaling pathways, which may be disturbed by HBV.

We then went on to check whether HBV-induced changes in the expression of some of the 35 DEGs might reciprocally affect HBV itself. We screened 7 of them and found only PGAM5 and PPP1R15A which could potently suppress all major HBV antigens expression in HBV recombinant cccDNA genome (**Fig.S6A-B)**. As one of the DEGs whose expression was downregulated by HBV (**Fig.2C, Fig.2E**), PGAM family member 5 (PGAM5) is a mitochondrial Serine/threonine-protein phosphatase that not only regulates the dynamics of mitochondria and the process of mitophagy but also is a central mediator for programmed necrosis induced by TNF or reactive oxygen species ^38^. Previously, PGAM5 deficiency was shown to protect acute liver injury driven by programmed necrosis^39^. The re-introduction of PGAM5 potently inhibited the expression of HBc, HBs and HBe (**Fig.2F-G**). HBV-induced down-regulation in PGAM5 expression may impact not only the dynamics and turnover of host mitochondria but also suppress the inflammation-induced necrosis and the tissue injury that could activate the host immune response.

PPP1R15A (Protein phosphatase 1 regulatory subunit 15A) could facilitate the recovery and survival of cells from stress. While the expression of PPP1R15A/GADD34 was up-regulated by HBV (**Fig.2C, Fig.2E**), re-introduction of PPP1R15A was found to significantly inhibit the expression of all three HBV antigens (**Fig.2F-G**).

To further dissect the mechanism of the suppressive effect of PPP1R15A and PGAM5 on HBV, co-immunoprecipitation coupled with mass spectrometry analysis was performed with empty vector as a negative control to identify deemed interactions between host proteins and PPP1R15A or PGAM5 (**Supplementary file S9**). The results shown that the host interaction proteins of PPP1R15A could participate in processes including metabolism of RNA, cellular response to stress, modulation by host of viral genome replication and regulation of translation (**Fig. S6C**); while PGAM5 could bind to host proteins involving metabolism of RNA, translation, mitotic cell cycle and mitochondrion organization (**Fig. S6D**). Additionally, to found viral target interact with PGAM5, we performed co-immunoprecipitation between PGAM5 and two proteins which were essential for virus replication, HBc and HBx. We found PGAM5 could bind to both of them, suggesting a possible perturbation of PGAM5 on HBV genome replication (**Fig. S6E**).

These data suggested that HBV infection and expression could impact on the proteostasis of many genes with meaningful functions and consequences on host-HBV interaction. Additionally, these observations have also testified the strength and power of dissecting host-HBV interaction using a multi-omics approach.

### HBV induced transcriptomic changes in host cells

To dissect host responses to HBV at transcription level, we further analyzed our RNA-seq data. To find potential host restriction factors of HBV cccDNA transcription, we performed gene ontology analysis and focused on GO term “transcription”, the heatmap of these genes was shown (**Fig. S7A-B**). An enriched GO term, “transcription corepressor activity”, caught our attention. The heatmap of 11 DEGs enriched in transcription corepressor activity was shown, and 2 genes, *SRSF2* and *SIRT6* were down-regulated by HBV (**Fig. 3A**). SRSF2 (serine and arginine rich splicing factor 2) is a component of spliceosome and responsible for pre-mRNA splicing and mRNA export from nucleus^40^. SIRT6 (Sirtuin 6) exhibits histone deacetylase activities that may participate in gene silencing^41^.

**Figure 3:**
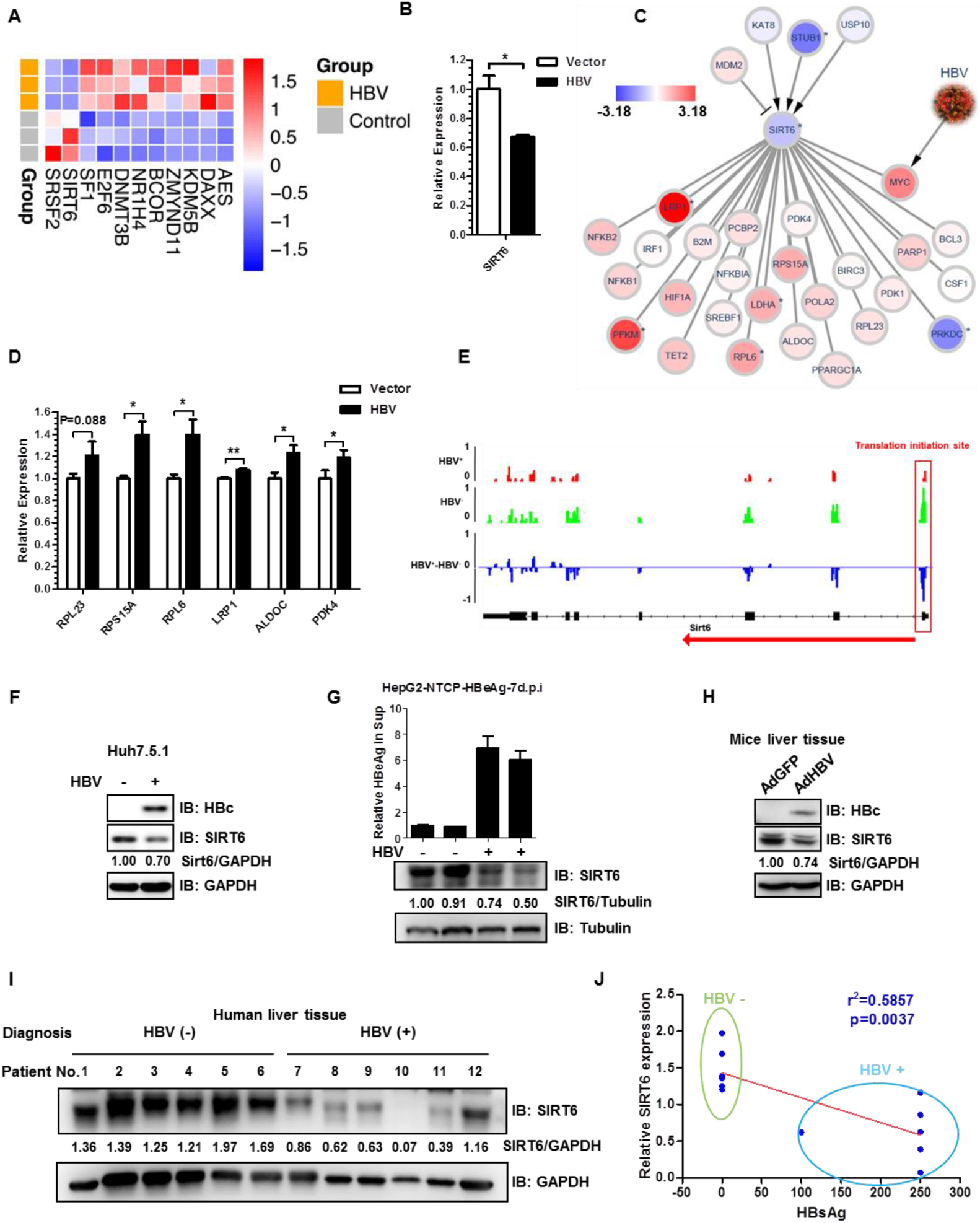
HBV induced transcriptional changes in host cells with SIRT6 emerged as a potential host virus-restricting factor. (**A**) RNA-sequencing was conducted with Huh7.5.1 cells transfected with Vector (Control) or prcccDNA and pCMV-Cre. Heatmap of 11 transcriptional DEGs, which have transcription corepressor activity, three biological replicates were shown for HBV and control group. (**B**) RT-qPCR confirmed that HBV down-regulated the transcriptional level of endogenous *SIRT6* transcription. (**C**) Network analysis of SIRT6-associated genes whose transcription was altered by HBV. Red and blue nodes indicate the up- and down-regulated genes, respectively. The color intensity indicates the fold change level of the gene. Nodes with * are DEGs (P-adjust <0.05, |log_2_FC|>=1, also see methods). (**D**) RT-qPCR validation of some of the SIRT6-associated genes that were shown in C. (**E**) Representative profiles of ribosome footprints of human SIRT6 ORF upon HBV, translation initiation of endogenous SIRT6 was down-regulated by HBV in Huh7.5.1 cells. (**F**) Endogenous *SIRT6* was down-regulated by HBV in Huh7.5.1 cells. (**G**) *De novo* infection of HBV down-regulated endogenous SIRT6 level in HepG2-NTCP cells, HepG2 cells stably expressing NTCP (sodium taurocholate cotransporting polypeptide), the functional receptor of HBV. The results of two independent biological replicates were shown. (**H**) HBV down-regulated SIRT6 in mouse livers infected with adenovirus harboring HBV genome. (**I**) Total proteins were extracted from the normal liver tissues of 12 patients who were diagnosed with HBV positive and negative, respectively. For patient information see **Table S1**. Endogenous SIRT6 or GAPDH proteins were visualized with IB using anti-SIRT6 or anti-GAPDH, with their relative abundances calculated using ImageJ. (**J**) The reverse correlation between the serum levels of HBsAg and the relative abundances of endogenous SIRT6 protein in liver tissues of the 12 patients. *, p<0.05, **, p<0.01, ***, p<0.001. qPCR results are presented as bar chart (For SIRT6, n=2; for other genes, n = 3); ELISA data are presented as bar chart (n = 3).

### Deacylase Sirt6 is down-regulated in patients tested positive of HBV antigens, or the cell and mouse models for HBV replication

*SIRT6* was previously established as an important regulator in controlling cellular response to stress, cellular component organization, carbohydrate metabolism and gene silencing, with histone H3 as its major targets ^42,43^. The transcription of endogenous *SIRT6* was significantly down-regulated by HBV as revealed by qRT-PCR analysis (**Fig. 3B**). To further characterize how HBV might also impact on SIRT6-related networks, we particularly looked into the transcriptional profiles of both the known upstream regulators and the downstream effectors of SIRT6 ^44–46^ (**Fig. 3C**). As shown in **Fig. 3C**, HBV down-regulated the transcription of *STUB1*, also known as CHIP, which prevents SIRT6 degradation through non-canonical ubiquitination^47^; while up-regulating the transcription of many genes involved in glucose or lipid metabolism such as *LRP1*, *PFKM* and *LDHA*, and that of ribosome protein genes such as *RPL6.* Notably, the transcription of MYC was also up-regulated in HBV-loaded cells^48^, which may also reflect the perturbation of SIRT6-related gene network. Some of the differences were further confirmed by qPCR analyses (**Fig. 3D**). Taken together, these results strongly suggested that SIRT6 could constitute a critical node mediating HBV-induced remodeling of host gene networks.

Consistent with RNA-seq results, the translation initiation of *SIRT6* was also compromised by HBV in RiboSeq (**Fig. 3E**). Immunoblotting analysis revealed that HBV did down-regulate the static level of endogenous SIRT6 protein in Huh7.5.1 cells (**Fig. 3F**), HepAD38 cells (**Fig. S7C**), or HepG2-NTCP cells for HBV infection (**Fig. 3G**). Moreover, in a recently developed mouse model for HBV persistence^25^, the level of endogenous SIRT6 protein was reduced in the HBV-infected (HBV^+^) mouse liver (**Fig. 3H, S7D**). To further examine the correlation between HBV and SIRT6 *in vivo*, total proteins were extracted from liver tissue samples of patients diagnosed with HBV positive or negative, respectively (detailed patient information was in **table S1)**. Indeed, the level of endogenous SIRT6 protein in these patients was negatively correlated to their serum HBsAg level **(Fig. 3I, 3J)**. Altogether, these data clearly indicated that HBV could target and down-regulate host SIRT6 *in vitro* and *in vivo*.

### Restoration of the homeostatic level of SIRT6 potently suppresses HBV gene expression

Subsequently, exogenous SIRT6, along with other members of the sirtuin family, was introduced into HBV^+^ Huh7.5.1 cells to test their potential effect on HBV gene expression, only SIRT6 strongly suppressed HBV gene expression (**Fig. S8A** and **S8B**).

Introduction of SIRT6-FLAG did markedly suppress HBcAg, HBsAg and HBeAg expression in multiple cell models for HBV infection and gene expression (**Fig. 4A-B, S8C, S8D**). Additionally, ectopic expression of SIRT6 also suppressed HBV gene expression in the context of both 1.1mer- and 1.3mer- HBV linear genome (**Fig.S8E-F**), while knockdown of endogenous SIRT6 with siRNA or shRNA elevated all HBV major antigens expression in HBV^+^ Huh7.5.1 and HepAD38 cells (**Fig.4C-D, S8G-H**), respectively. Furthermore, the deacetylation activity of SIRT6 seemed to be essential for its restrictive effect on HBV, as its HBV-suppressing effect was largely abolished by the point mutation S56Y, G60A, R65A or H133Y that disrupted the deacetylation activity of SIRT6^49^ **(Fig. 4E-F)**, Southern blotting analysis confirmed that SIRT6 could suppress HBV genome replication in both HepG2 (**Fig.4G**) and Huh7 cells (**Fig.S8I**), while SIRT6 containing mutation S56Y only had marginal effect (**Fig.4G**). Therefore, endogenous SIRT6 was emerging as a novel host virus restriction factor (Vrf) and restoration of SIRT6 homeostasis could potently suppress HBV gene expression and genome replication.

**Figure 4:**
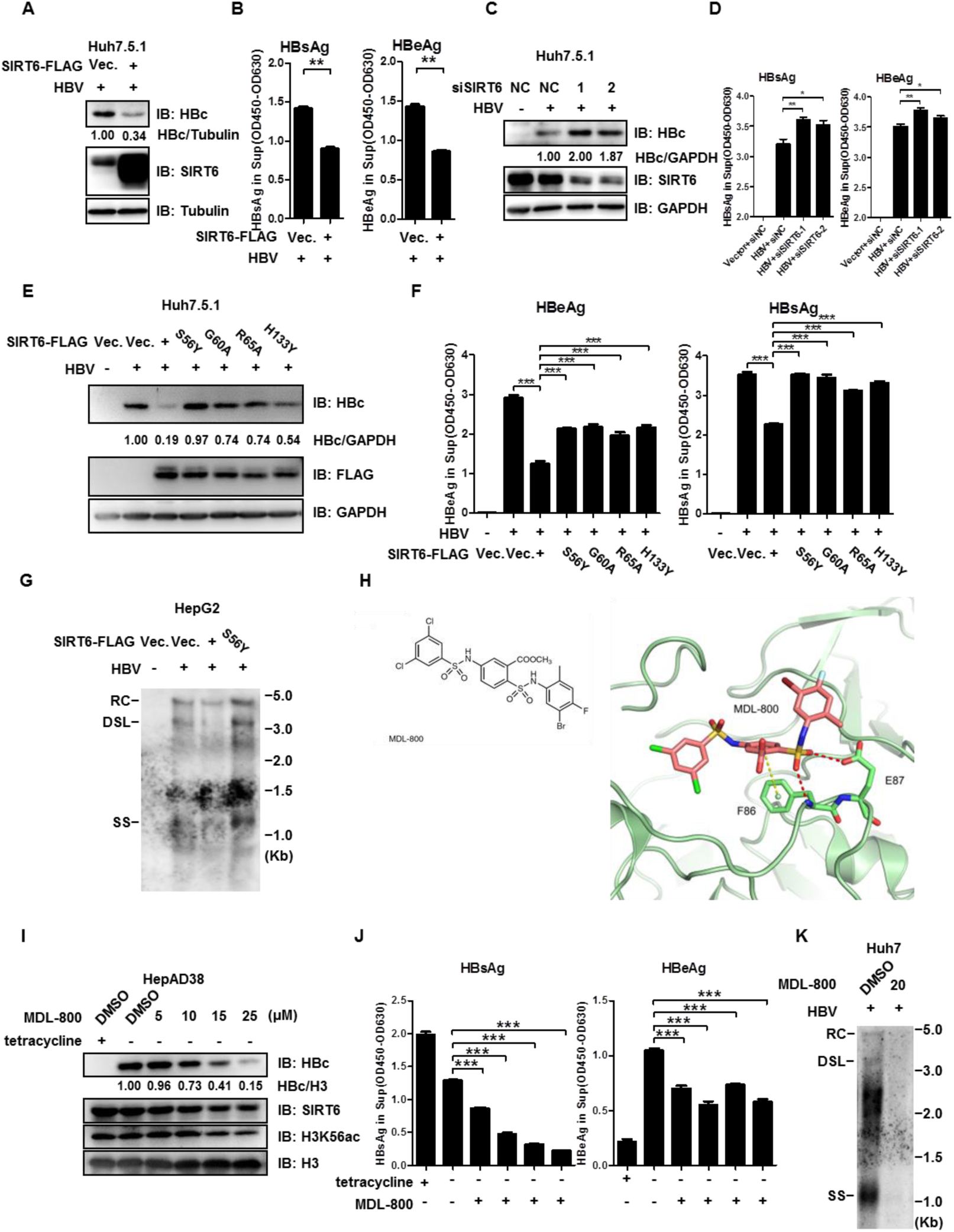
Identification of SIRT6 as a host virus-restricting factor for HBV. (**A** and **B**) Introduction of Sirt6 suppressed the expression of HBc (A) and HBs and HBe (B) antigens in Huh7.5.1 cells. (**C** and **D**) Knocking-down endogenous Sirt6 promoted the expression of HBc (C) and HBs and HBe (D) antigens in Huh7.5.1 cells. (**E** and **F**) Effect of SIRT6 enzymatic mutants on HBc (E) and HBs and HBe (F) expression in huh7.5.1 cells. (**G**) The effect of SIRT6 wild type or enzymatic deficient mutant on HBV genome replication was measured via southern blotting in HepG2 cells. (**H**) The chemical formula and a close-up view of MDL-800 in complex with 2′-O-acyl-ADP ribose (2′-O-acyl-ADPR) motif of human SIRT6 (X-ray structure, PDB number 5Y2F). ELISA shows that MDL800 suppressed the expression of HBc (**I**) and HBs and HBe (**J**) antigens in dose-dependent manner in Tet-controlled HBV in HepAD38 cells. (**K**) MDL-800 treatment significantly suppressed HBV genome replication intermediates in Huh7 cells. RC, relaxed circular DNA; DSL, double strand linear DNA; SS, single strand DNA. For ELISA, for panel B, n=2; For others, n=3. *, p<0.05, **, p<0.01, ***, p<0.001.

Recently, MDL800 was developed as a specific allosteric activator for human SIRT6^22^ (**Fig.4H**). In HBV mini-chromosome, H3K56ac is an epigenetic marker for active transcription of HBV cccDNA^3^, whose acetylation status is dynamically controlled by histone acetyl transferase CBP/p300^50^ and deacetylase SIRT6^43^. In HepAD38 cells (**Fig.4I-J**), HBV^+^ Huh7.5.1 (**Fig. S9A, S9B**) or HepG2 (**Fig. S9C, S9D**) cells, the levels of HBcAg, HBsAg and HBeAg, as well as H3K56ac were decreased upon MDL800 treatment in a dose-dependent manner. Moreover, the suppressive effect of MDL800 on HBV gene expression was largely abolished when knocking down the endogenous SIRT6 in HepG2 cells (**Fig. S9E**). Furthermore, Southern blotting analyses clearly indicated that MDL800 treatment also potently suppressed HBV genome replication (**Fig. 4K**). These data suggesting that SIRT6 could suppress HBV gene expression and genome replication via de-acetylating H3K56ac, regardless of host cell types.

### SIRT6 suppresses rcccDNA transcription involving interacting with HBcAg and direct deacetylating of histone H3 in HBV mini-chromosome

To test which proteins in sirtuins family could interact with HBc, co-IP assay was performed and the result shown that only SIRT6 could strongly bind to HBc, and SIRT7 has a week interaction with HBc, while others couldn’t (**Fig. S10A**). To further investigate how SIRT6 mediated HBV restriction, co-IP assay was performed between SIRT6 family and HBcAg mutually (**Fig. 5A**). HBcAg, but not HBx, was found to interact with SIRT6 involving the core domain (**Fig.S10B-C**). Immunofluorescence assay further confirmed that SIRT6 and HBcAg indeed co-localized within speckles in nucleus during HBV replication (**Fig. 5B**). ChIP assay with SIRT6 antibody indicated that endogenous SIRT6 could directly bind to mini-chromosome during HBV replication with LINE1 as a positive control^51^ (**Fig. 5C**). HBcAg protein is known to be important for formation and maintenance of HBV mini-chromosome, promoting the epigenetic permissive state for HBV *in vivo*^3,52,53^. The interaction between HBcAg and SIRT6 might help recruit SIRT6 to HBV mini-chromosome and deacetylate the histones for transcriptional repression. Another ChIP-qPCR assay using H3K56ac (acetylation on histone H3 lysine 56) or H3K9ac (acetylation on histone H3 lysine 9) antibodies revealed that overexpression of SIRT6 alone was sufficient to deacetylate histone H3 on HBV mini-chromosome (**Fig. 5D**), and such effect could be reversed when endogenous SIRT6 was knocked-down (**Fig. 5E**). Moreover, MDL800 treatment alone was sufficient to deacetylate histone H3K56ac on HBV mini-chromosome (**Fig. 5F**). Therefore, SIRT6 appeared to suppress HBV gene expression through epigenetically silencing the virus mini-chromosome.

**Figure 5:**
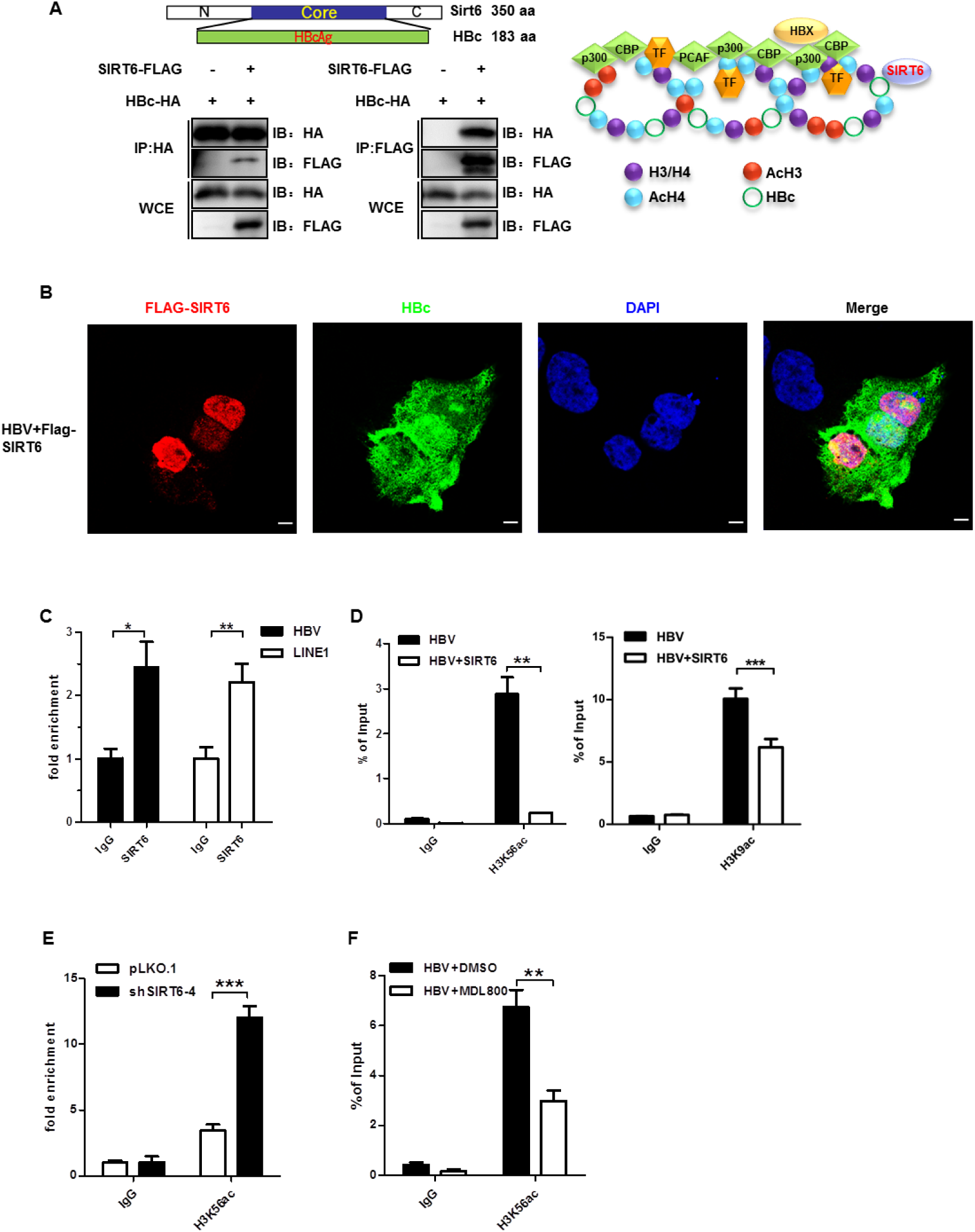
SIRT6 restricts HBV gene expression through deacetylating H3K9ac and H3K56ac on HBV mini-chromosome. (**A**) SIRT6 co-immunoprecipitated with HBcAg in 293FT cells. (**B**) SIRT6 and HBcAg co-localized in the nuclei of Huh7.5.1 cells during HBV replication. Scale bars, 20 μm. ChIP assays were performed using indicated antibodies with cells upon either HBV transfection alone (**C**), or in combination with SIRT6 overexpression (**D**) or knockdown (**E**) or MDL800 treatment (**F**). *, p<0.05, **, p<0.01, ***, p<0.001. ChIP data were acquired in Huh7.5.1 cells and were presented as bar chart (n=3 per group).

### MDL800 suppresses HBV gene expression *in vitro* and *in vivo*

To further access the effect of endogenous SIRT6 on HBV infection and virus gene expression in *de novo* cell infection and animal system. MDL800 was tested in HepG2-NTCP cells^24^ and a recently established mouse model of HBV through tail-vein hydrodynamic injection (HDI)^25^, respectively. Firstly, MDL800 potently suppressed HBe and HBs expression in *de novo* infection in HepG2-NTCP cells at varying time points (**Fig. 6A-B**). A group of mice were subjected to intraperitoneal injection with vehicle only or MDL800 continuously for two weeks after intravenous injection of rcccDNA system of HBV (see methods section). As shown in **Fig.6C**, continuous administration of MDL800 did lead to significant reduction in serum levels of HBsAg at different time points, without elevating serum ALT (alanine aminotransferase) activities (**Fig.6D**) or causing obvious morphological damage in the liver tissues (**Fig. 6E**). Immuno-histochemical staining demonstrated that MDL800 did lower HBcAg expression and the level of H3K56 acetylation in mouse hepatocytes (**Fig. 6E-F**). Taken together, these results demonstrated that MDL800 could suppress HBV gene expression by specifically augmenting the de-acetylase activity of SIRT6. MDL800 thus appeared to be a promising lead compound for future HBV treatment, through both lowering HBV DNA loads and silencing virus gene expression.

**Figure 6:**
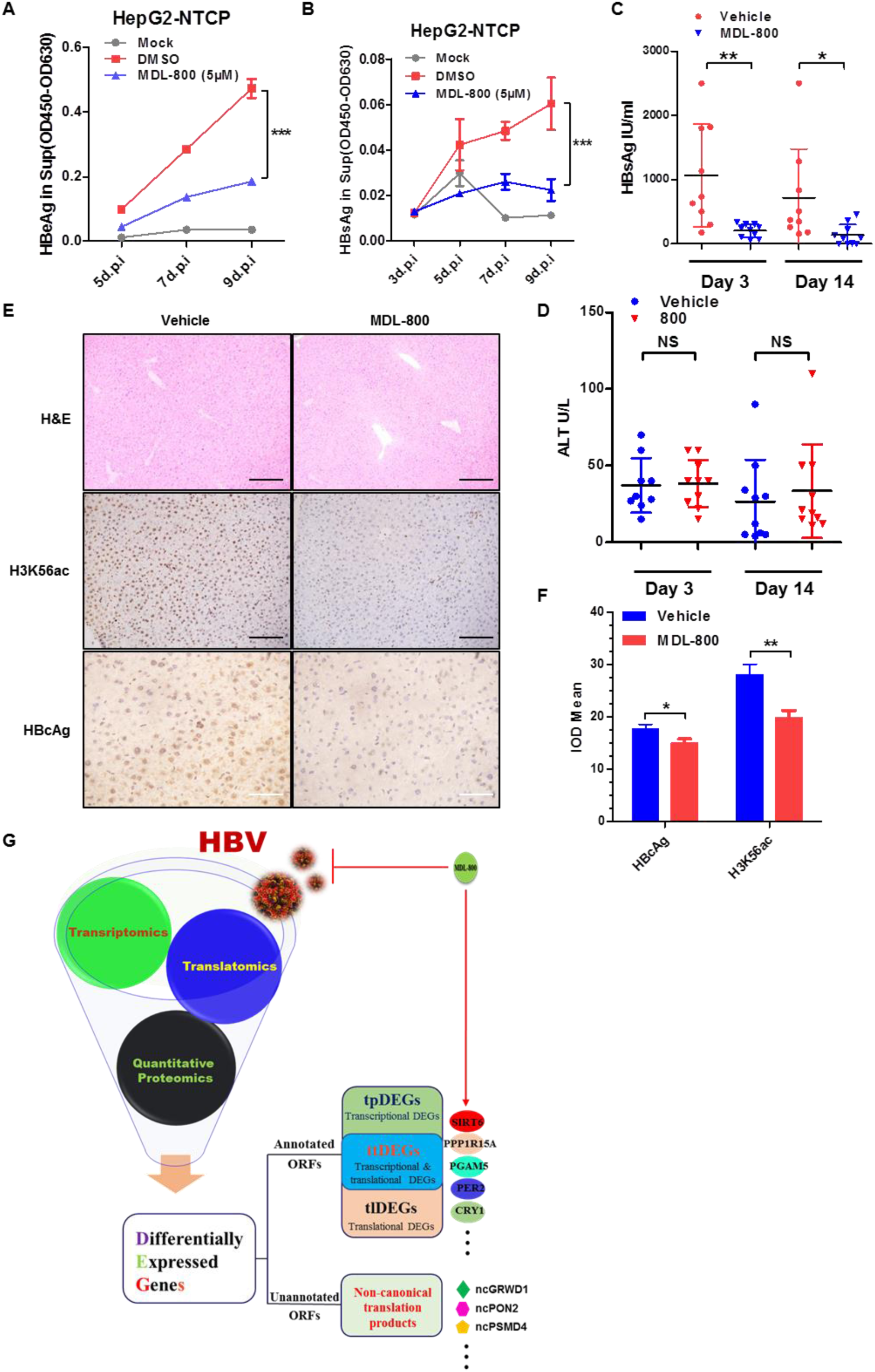
MDL-800 restricts HBV *in vitro* and *in vivo*. (**A** and **B**) HepG2-NTCP cells were infected either with vehicle (mock) or HBV particles purified from the culture medium of HepAD38 cells, the HBV groups were treated with DMSO or MDL800, and the HBeAg (**A**) and HBsAg (**B**) level in culture medium of each group were collected and measured at indicated time points. (**C**) A Mouse model for HBV infection was established as previously described^25^. Peripheral blood samples were collected and subjected to ELISA to detect serum HBsAg, for vehicle group and MDL800 group. (**D**) Peripheral blood samples of mice treated with MDL800 or vehicle were collected and subjected to Alanine-Aminotransferase (ALT) assay. (**E**) HE or immunohistochemistry staining of liver sections from mice receiving vehicle or MD800 from experiment in F. Representative images of indicated group were shown. For the panel labeled H&E and H3K56ac, Scale bars, 200 μm; For the panel labeled HBcAg, Scale bars, 100 μm. (**F**) Statistics analysis of IHC results of indicated group, the mean integral optical density (IOD mean) of five random visual fields for each sample (n=9 for vehicle group and n=10 for MDL800 group) was measured. (**G**) Summary of the major findings in this study. Multi-omics interrogation into HBV-host interaction has led to the discovery of multiple HBV-induced DEGs including canonical and non-canonical genes in host cells. For panel A and B, two-way ANOVA test was used, for others, student t test was used. *, p<0.05, **, p<0.01,***, p<0.001.

## Discussion

HBV represents a yet unresolved global threat to public health, and a leading cause of mortality worldwide. Despite the luminating goal by WHO’s worldwide campaign to eliminate HBV infection from the top list of health threat in 2030, complete curing treatment is still unavailable and more “functional treatment” yet to be developed, mostly due to our limited understanding of HBV-host interaction and HBV life cycle ^1,5^. Application of the state-of-art-omics techniques were thus called upon to fully elucidate the mechanism of cccDNA transcription and identify novel targets that would prove critical for reaching a functional cure^1,54^. To this end, we have applied RiboSeq, SILAC and RNA-seq, techniques to globally interrogate HBV-host interaction.

A total of 20,533 non-canonical ORFs were identified by ribosome profiling (**Fig. 1B**), which generally encoded proteins or polypeptides shorter than those encoded by the canonical ORFs (cORFs) (**Fig. 1C**). As a significant portion of the ncORFs overlapped with many annotated ORFs that encoded proteins of known functions, it was convenient to speculate that altered translation of these ncORFs might affect translation of the concerned cORFs, and very likely change the homeostasis and functionality of the canonical proteins. Particularly, the production of *ncPON2* was down-regulated by HBV, and ectopic expression of ncPON2 suppressed viral gene expression (**Fig. 1D** and **1J**), this phenomenon suggested that ncPON2 was a novel viral restriction factor of HBV. As little was known about the functions of the translational products of these novel ncORFs, our work combining RiboSeq and SILAC has offered an opportunity to investigate the roles and dynamics of the translational products of ncORFs during HBV-host interaction. On the other hand, our findings also revealed a previously unknown effect of HBV infection on the translational plasticity of host genome.

There has been much advance in generating renewable sources of hepatocyte-like cells from primary hepatocytes^55^. Currently, heterogeneity as well as the low transfection and/or infection efficiency of the newly transformed cells has limited their application as stable models for HBV infection^13^. Therefore, several classic cell-based models for HBV study were employed in this work for screening host restriction factors and primarily confirming the findings, namely Huh7.5.1, Huh7, HepG2 and HepAD38, mainly due to their extraordinary reproducibility and robustness for supporting hepatitis B virus replication and gene expression ^56–58^. RNA-seq analysis revealed that HBV altered the transcription of 533 previously annotated host genes, which defined the set of HBV-specific transcriptional differentially expressed genes (tpDEGs) (**Fig S5B, supplementary file S6**). Through reprograming the transcription of these tpDEGs that were highly enriched in key pathways from protein synthesis and processing to RNA splicing and to amino acid and central carbon metabolism (**Fig.S5C**), HBV might reshape the homeostasis and functionality of these pathways to manipulate the activities of the concerned processes.

At the level of translation, RiboSeq revealed that HBV induced a major shutoff on mRNA translation of 939 previously annotated host genes, seemingly to favor the expression of virus genes, while specifically up-regulating the translation of 39 host genes (**Fig S5B, supplementary file S7**). In KEGG enrichment analyses, these 978 translational DEGs (tlDEGs) were typically over-presented with several signaling pathways from necroptosis to Hippo signaling, which might profoundly impact the cellular responses to HBV (**Fig S5D**).

Interestingly, a small set of ttDEGs whose expression was impacted by HBV both transcriptionally and translationally (**Fig.2C**). Importantly, HBV gene expression seemed to be pronouncedly affected upon the reinstatement of the homeostasis of SIRT6, and PGAM5, suggesting that SIRT6 and PGAM5 might serve as host-restrictive factors (**Fig.2F-G, Fig.4A**). On the other hand, stress-inducible protein PPP1R15A was up-regulated by HBV and its ectopic expression was found to suppress HBV. Given the protective role of PPP1R15A in preventing liver injury, did HBV up-regulate PPP1R15A to put on a self-restriction on its own proliferation and gene expression while suppressing the host immune response? This question remains to be answered.

Interestingly, a small set of genes which has transcription corepressor activity was impacted by HBV transcriptionally (**Fig. 3A**), and SIRT6 may represent one of the critical epigenetic regulating nodes targeted by HBV. Indeed, the expression of upstream regulators and down-stream effectors mediated by SIRT6 were altered by HBV (**Fig. 3C** and **3D**). Importantly, HBV was found to down-regulate endogenous SIRT6, not only in the cell-based HBV infection models, but also in HBV infected patient’s livers and HBV-loaded mice (**Fig. 3E-I, Fig. S7C** and **S7D**). Moreover, HBV gene expression seemed to be pronouncedly affected upon the changed level of SIRT6, suggesting that SIRT6 might serve as host-restrictive factors (**Fig. 4A, S8C, S8D**). Mechanistically, overexpression of SIRT6 was found to curtail HBV DNA replication and silence virus gene expression involving sequestering HBcAg during viral replication and deacetylating Histone 3 at K9 and K56 through directly binding to mini-chromosome (**Fig. 5A-E**).

In principle, targeted manipulation of the expression and functionality of HBV-specific ttDEGs may thus represent unprecedented opportunities to combat HBV infection and the related diseases. Recent development of MDL-800 as a specific allosteric agonist of SIRT6 ^22^ has empowered us to test whether chemical activation, rather than overexpression, of endogenous SIRT6, would have any effect on HBV replication and gene expression. The structure of MDL800 and SIRT6 co-crystallization was determined before and the specificity and effectiveness of MDL800 was confirmed in SIRT6 knockout hepatoma cell lines and by using *in vitro* synthesized KQTARK-ac-STGGWW peptide, respectively^22^. With the encouraging data that MDL-800 could efficiently suppress HBV gene expression and/or genome replication in transient transfection cell systems, stable replication cell systems, *de novo* infection cell systems, and mouse models with negligible toxicity (**Fig. 4I-K, 5F, 6A-B, 6C-F, Fig. S9A-D**), MDL-800 was thus emerging as a promising lead compound for HBV treatment with clear and specific mode of action.

Altogether, we have presented here a multiomics landscape of the HBV-host interaction through ribosome profiling, SILAC and RNA-sequencing analysis. By identifying translational products of the ncORFs and discovering multiple transcriptional DEGs, which all contributed to HBV-induced changes in the proteostasis network of host cells, these findings have opened an new avenue to identify potential drug targets, biomarkers, neoantigens or even lead compound, as showcased with ncGRWD1, ncPON2, PPP1R15A, PGAM5, SIRT6, and MDL-800 (**Fig.6G**). To develop novel therapeutics, and potential diagnostics and prognostics in combating microbial infection, more -omics work should be performed to gain better global and detailed view of virus-host interactions^1^. This study was but one example.

## Materials and Methods

### Cells, antibodies, reagents and constructs

Huh7.5.1, HepG2, HEK293T, HEK293FT and HepG2-NTCP cells were maintained in Dulbecco’s modified Eagle’s medium (DMEM) (Hyclone) supplemented with 10% fetal bovine serum (FBS) and 100 units/ml of penicillin-streptomycin (Gibco), in a humidified incubator supplemented with 5% CO2 at 37°C (Thermo Scientific). HepAD38 cells were cultured as previously described^26^. (For details of antibodies, reagents and constructs, see **Supplementary file S11**).

### RiboCode analysis and annotation

Here, non-rRNA sequencing reads were aligned to human genomic reference (hg38) using STAR program. Then, the Ribocode^29^ pipeline was used to determine translated regions in ribosome profiling data from HBV and control group.

ORFs were annotated according to the location of their start codons in the structure of the original ORFs: “annotated” (are perfectly matched to the previously known annotated ORFs); “unanotated” (are translated from RNAs from the “non-transcribed” intergenic or intragenic regions in human genome); “nonoverlap_uORF” (with start codons upstream to those of the previously annotated ORFs, with the resulted frames not overlapping with the known ORFs); “nonoverlap_dORF” (with start codons downstream to those of the annotated ORFs, not overlapping with the annotated ORFs), “Overlap_uORF” (with start codons upstream to those of the previously annotated ORFs, with resulted frames overlapping the annotated ORFs), “Overlap_dORF” (with start codons downstream to those of the annotated ORFs, overlapping to the annotated ORFs), “Internal” (Start codons in the internal region of the known ORFs, with resulted frames not overlapping with those of the original ORFs); “Other” (ORFs that originated from RNAs transcribed either from the intergenic genomic regions defined as “non-coding” before, or from the on-transcribed regions in the known genes).

### Stable isotope labeling by amino acids in cell culture (SILAC)

SILAC was performed as previous described, briefly, HepG2 or HepAD38 cell lines were passaged at 80% confluence in heavy (L - lysine - 2HCL(13C6 15N2, 98% isotopic purity) and L –arginine - HCL(13C6 15N4, 98%)), middle (L - lysine - 2HCL (4,4,5,5 D4, 98%) and L –arginine - HCL(13C6, 98%)) or light (normal) SILAC media. Cells were grown to confluence and passaged for ten passages. Cells were collected at the 4^th^, 8^th^, or 10^th^ for determination of labeling efficiency, when it reached 95%, cells were washed with ice-cold PBS, and then lysed with SDT buffer (2% SDS(w/v), 0.1M DTT, 0.1M Tris, pH 7.6), cell lysates were collected and boiled, and centrifuged at 15000g for 15 minutes, the supernatants were collected and stored at −80°C. Before mass spectrometry analysis, Proteins in each cell lysate were quantitated by gel-electrophoresis and Commassie brilliant blue staining, and imagJ analysis, and 20μg from heavy, middle and light conditions were mixed together and then subjected to mass spectrometry at Beijing Proteome Research Center, BPRC. Data were searched using the Maxquant search engine^31^ with two sequence databases, respectively, including normal human protein sequence from UNIPROT database and protein sequence of non-canonical ORFs from RiboCode. Manual analysis of MS/MS matches confirmed 47 ncORF peptide sequences (**Supplementary File S3**).

### Ribosome profiling, RNA-seq, and data processing

Ribosome profiling was performed as previously described^59^, with some modification. Briefly, before cell lysis, the control groups and experiment groups were pretreated with Harringtonine at the concentration of 2μg/ml in 37°C for 120s, and then add cycloheximide at the concentration of 100μg/ml, mix well and proceed to the next step quickly. Lysates of a 10 cm dish cells were treated with 750 U RNase I (Invitrogen, cat. No. AM2294) for 45 minutes at room temperature, then transferred to an ultracentrifuge tube on a sucrose cushion (∼34%) and centrifuged at 70,000 rpm at 4°C for 4 hr. (Hitachi CS150GX). Ribosome-protected fragments were purified using miRNeasy RNA isolation kit (QIAGEN, cat. No. 217004). RNA was size selected, and then dephosphorylated, linker ligated, and then subjected to RT-PCR, circularization, rRNA depletion, and 10-12 cycles of PCR. The enzymes used were T4 PNK (NEB, cat. No. M0201S), T4 RNA ligase 2, truncated (NEB, cat. No. M0242S), Universal miRNA Cloning Linker (NEB, cat. No. S1315S), Superscript III (Invitrogen, cat. No 18080-044), Circ Ligase (Epicentre, cat. No. CL4111K), NEBNext® High-Fidelity 2X PCR Master Mix (cat. No. M0541L). Resulting fragments were size selected from an 8% acrylamide non-denaturing gel and purified by incubation with DNA gel extraction buffer (300 mM NaCl, 10 mM Tris (pH 8) and 1 mM EDTA). Ribosome footprint libraries were analyzed on agilent 2100 bioanalyzer and sequenced on HiSeq2500 platform. The sequencing data was preprocessed by discarding low-quality reads, trimming adapter sequence, removing ribosomal RNA (rRNA) derived reads^59^. Next, non-rRNA sequencing reads was aligned to human genomic reference (hg38) using HISAT2. The abundance of these transcripts in each sample was computed with StringTie and Ballgown^60^, only genes with reads numbers above 5 were selected. Differential expression was determined using one-side T-test. Genes with p-value <=0.05 and |FC| >=1.5 were considered as differentially expressed genes (DEGs).

For RNA-seq, total RNA of cells was extracted by Trizol (Invitrogen) according to manufacturer’s instructions, and then poly-A-selected using NEBNext® Poly(A) mRNA Magnetic Isolation Module (Catalog # E7490S). The Quality of the poly-A-selected RNA was analyzed using Agilent 2100 Bioanalyzer. Library preparation using Illumina TrueSeq mRNA sample preparation kit (Catalog IDs: RS-122-2001) was accomplished at the National Center of Plant Gene Research (Shanghai), and cDNA library was sequenced on Illumina HiSeq 2500. The sequencing reads was aligned to human genomic reference (hg38) using HISAT2. The abundance of these transcripts in each sample was computed with StringTie and Ballgown^60^. Differential expression was determined using DESeq2. Genes with adjust p-value <=0.05 and |FC| >=1.5 were considered as differentially expressed genes (DEGs). Gene Ontology enrichment of the identified DEGs was performed using DAVID. The GO terms of transcription molecular function were selected. Volcano plots, heatmap were drawn in RStudio with the ggplot2 packages. The visualization of mapping results was performed by Integrative Genomics Viewer^61^.

### Western blotting

Cells were lysed in 1× SDS-PAGE loading buffer (10% glycerol, 50 mM Tris-HCl, pH 6.8, 1% β-mercaptoethanol, 0.08% bromophenol blue, 2% SDS). Protein electrophoresis was conducted using SDS-PAGE gels, transferred to PVDF membranes (Bio-Rad). Membranes were blocked, incubated with primary antibodies overnight at 4°C, and then washed with TBST for three times, incubated with secondary antibodies for 60 min at room temperature, and again washed in TBST for three times, and then incubated with high-sig ECL western blotting luminol/enhancer solution (Tanon, Cat. No. 180-5001). Protein bands were visualized using Tanon-5200 (Tanon). The bands intensity was calculated by ImagJ.

### Cell transfection

The transfection of Huh7.5.1 cells and Huh7 cells in ribosome profiling and RNA-seq experiments were transfected with indicated plasmids using Lipofectamine 2000 (Life Technologies). si-RNA transfection was performed using MaxFection™ 8600 Transfection Reagent (Biomaterials USA, cat. No. MF8600-001) according to the manufacturer’s instructions. Other transfection experiments were conducted using polyethylenimine (Sigma) if not otherwise mentioned. (For the sequences of siSIRT6, see **Supplementary file S11**).

### Stable gene knockdown and overexpression

Lentiviral particles harboring SIRT6 overexpression vector (pCDH; Addgene) or *sirt6* shRNA expression vector (pLKO.1; Sigma-Aldrich) were produced by transfection of HEK293FT cells with indicated plasmids and lentiviral packaging plasmid mix. HepG2 or HepAD38 cells were transduced by the harvested viral supernatants in the presence of 8 μg/ml polybrene (Sigma), followed by selection with 2 μg/ml puromycin (Clontech). (For the sequences of shSIRT6, see **Supplementary file S11**).

### Co-immunoprecipitation (Co-IP) assay

For Co-immunoprecipitation assay, 48h after transfection, HEK293T cells were incubated in co-IP buffer (50 mM Tris-HCl, pH 7.4, 150 mM NaCl, 5mM EDTA, 10% Glycerol, 0.5% NP-40) plus 1 mM NaF, 1mM Na_3_VO_4_ and 1% protease inhibitor cocktail (bimake, cat. No. B14001), followed by ultra-sonication. After spin at full speed at 4°C for 10min, the corresponding antibody-conjugated beads were added into supernatant. After incubation at 4°C, the beads were washed and boiled in 2× SDS-PAGE loading buffer, and then subjected to western blotting with indicated primary antibodies.

### Immunofluorescence assay

For immunofluorescence assay, cells were washed with phosphate buffered saline, fixed with 4% paraformaldehyde, then permeabilized for with 0.1% Triton X-100, and then blocked at room temperature for 1h in 1.0% BSA, after wash with PBST, cells were incubated with indicated primary antibodies, and then incubated with Alexa Fluor 488 (A11029, Thermo Fisher) or Alexa Fluor 647 (A21245, Thermo Fisher)-conjugated secondary antibodies. Cell nucleus was stained with DAPI (Thermo Fisher). Images were obtained using an Olympus BX51 microscope (Olympus) or Leica TCS SP8 confocal microscope (Leica).

### ELISA

HBsAg and HBeAg from supernatants of HBV replicating cells were measured using the ELISA kits (Shanghai Kehua Bio-engineering Co., Ltd) according to the manufacturer’s instructions. The medium was changed the day before collection.

### Chromatin immunoprecipitation (ChIP)

ChIP for H3K56ac, H3K9ac and SIRT6 was performed according to the standard protocols^23^, with some modifications. Briefly, hepatoma cells were suspended and cross-linked with 1% formaldehyde at room temperature for, quenched with 2.5 M glycine, washed in ice-cold PBS buffer, and lysed with 1×SDS lysis buffer (1% SDS, 10 mM EDTA, 50 mM Tris-HCl, pH 8.0). Cellular lysates were sonicated with high power, 30s on 30s off for 35 cycles, diluted in ChIP dilution buffer (SDS 0.01%, 1.1% Triton X-100, 1.2 mM EDTA, 16.7 mM Tris-HCl, 167 mM NaCl, pH8.0), and immunoprecipitated with indicated antibodies. Normal rabbit IgG (Santa Cruz biotechnology) was used as negative control. Immunoprecipitates were collected with Protein A/G-agarose beads (Merk millipore) and washed sequentially with low-salt wash buffer (0.1% SDS, 1% Triton X-100, 2 mM EDTA, 20 mM Tris-HCl, pH 8.0, 150 mM NaCl), high-salt wash buffer (0.1% SDS, 1% Triton X-100, 2 mM EDTA, 20 mM Tris-HCl, pH 8.0, 500 mM NaCl), LiCl immune complex wash buffer (0.25 M LiCl, 1% NP40, 1% deoxycholate, 1 mM EDTA, 10 mM Tris-HCl, pH 8.0), and TE buffer (10 mM Tris-HCl, 1 mM EDTA, pH 8.0), DNA–protein immune complexes were eluted by elution buffer (1% SDS, 0.1 M NaHCO_3_). Then NaCl were added and samples were heated to 65 °C for 4 hours, and then treated with proteinase K at 45 °C for 1 h. DNA was purified by standard phenol chloroform extraction protocol and assayed by quantitative PCR on a CFX96 real-time PCR system (Bio-Rad) or LC96 (Roche) with Talent qPCR PreMix (SYBR Green, TIANGEN). Fold enrichment was calculated as percentage input and normalized to total H3. (For the sequence of qPCR primers, see **Supplementary file S11**).

### Southern blot analysis of HBV DNA

HBV DNA Southern blot was conducted following a modified procedure as described previously ^62^. Briefly, after transfected with HBV rcccDNA, Huh7.5.1 cells were lysed with 0.5% NP-40 in TBS (10 mM Tris-HCl [pH 7.0], 150 mM NaCl), Nuclei were pelleted by brief centrifugation. To selectively extract HBV DNA from intracellular core particles, cytoplasmic lysates were treated with micrococcal nuclease (Amersham Biosciences) to remove input plasmid DNA. After 1% SDS digestion containing protease K for 2 h at 55°C, viral DNA was ethanol precipitated. rcccDNA in the nuclei was purified with similar procedure in Hirt extraction^63^. The nuclear pellet was resuspended in 1 ml SDS lysis buffer (50 mM Tris-HCl [pH 8.0], 10 mM EDTA, 150 mM NaCl, and 0.5% SDS), mixed with 0.25 ml of 2.5 M KCl, and incubated at 4°C with gentle rotation overnight. The lysate was centrifuged at 14,000 ×g for 20 min and rcccDNA was further extracted with phenol and chloroform, followed by ethanol precipitation.

### Purification of HBV virus

HepAD38 cells were maintained in complete DMEM/F-12 medium in the presence of 2% DMSO. After tetracycline withdraws for 12 days, the culture medium was filtered through 0.45μm filter and then precipitated at 4°C in the presence of 8% PEG8000 overnight. The pellets were collected through centrifugation at 10000g and suspended in PBS with 20% glycerol, after brief centrifugation, the aliquots were stored at −80°.

### HBV *de novo* infection of HepG2-NTCP

HBV infection experiment was performed as described previously^24^, with some modifications. Briefly, HepG2-NTCP cells were seed in 24-well plates with complete DMEM for 24 h, and then cultured with PMM medium for another 24 hours. The cells were then inoculated with HBV purified from culture medium of HepAD38 cells at multiplicity of 100 genome equivalents for 16∼24 hours at 37°C, then each well was washed with 500 microliter PMM for three times, and then 500 μl fresh PMM were added in the presence or absence of 5μM MDL800. Mock infection was inoculated with PMM medium in the presence of 4% PEG8000. Cells were maintained in PMM, and medium was changing every other day until supernatants were collected.

### Mice and *in vivo* chemical test

Mouse study was conducted as described previously^25^. Briefly, wild type (WT) male mice (C57BL/6) (4-5 weeks) were hydrodynamic injected with a modified rcccDNA system harboring *β*2-microglobulin-specific shRNA (shB2M) to reduce T-cell response. 4 μg prcccDNA-shB2M and 4 μg pCMV-Cre were co-injected within 5 to 8s through tail veins in a volume of DPBS equivalent to 8% of the mouse body weight. Alb-Cre Transgenic mice (C57BL/6-Tg [Alb-Cre] 21Mgn/J) using albumin promoter to express Cre recombinase were obtained from Jackson Laboratory (Bar Harbor, ME). For Ad-GFP/rcccDNA transduction, 1.5 ×10^9^ PFU of vehicle were intravenously introduced into Alb-Cre Transgenic mice (6–8 weeks).

For *in vivo* test of MDL800, mice were randomly divided into two groups according to the HBsAg unit in serum to avoid bias 3 days after plasmid co-injection; MDL800 was dissolved in 5% DMSO, 30% PEG-400, 65% saline, with 1.5 Meq 1N NaOH, and administrated via intraperitoneal injection at the dose of 65mg/kg body weight/day continuously for two weeks. The investigators were not blinded to the group allocation. All animal studies were approved by the Animal Ethics Committee of Institute Pasteur of Shanghai (no. A2012008-2), Chinese Academy of Sciences.

### *In vitro* chemical test

MDL800 was resolved in DMSO (Sigma) with 100mM. For Huh7.5.1 and HepG2 cells, MDL800 was added with indicated final concentrations by changing fresh medium at day 1 and 2 post prcccDNA and pCMV-Cre transfection. The supernatants and cell lysates were harvested at day three post transfection and subjected to ELISA and western blotting. For HepAD38, after tetracycline removing, chemical was added similar to Huh7.5.1.

### Patients

Liver biopsies were acquired from Ruijin Hospital and Eastern Hepatobiliary Surgery Hospital and stored at −80° C before use. This study was approved by the Institutional Ethics Review Committee at Ruijin Hospital and Ethic Committee of Eastern Hepatobiliary Surgery Hospital. Written informed consent was obtained from each patient.

### Statistics

All experimental data were expressed as ± standard deviation (SD). Unpaired Student’s two-tailed *t*-test was performed with GraphPad Prism software. All experiments were performed at least three times independently, only *P* value of *<* 0.05 was considered to be statistically significant, *P* <0.05, *; *P* <0.01, **; *P* <0.001, ***.

## Supporting information

Supplementary file S2. Ribocode analysis

Supplementary file S3. Novel peptides

Supplementary file S4. ncGRWD1 ncPON2 interactor

Supplementary file S5. FPKM of RNAseq

Supplementary file S6. DEGs list of RNAseq

Supplementary file S7. DEGs list of ribosome profiling

Supplementary file S8. Translation_efficiency

Supplementary file S9. PPP1R15A PGAM5 interactor

Supplementary file S10. proteome

Supplementary file S11. List of resources used in this study

Supplementary file S1. FPKM of ribosome profiling

## Acknowledgments

We specifically acknowledge the excellent support from Dr. Chao Peng and the proteomics facility at the National Center for Protein Science (NCPS) Shanghai and the molecular biology, and cell biology in SIBCB. We thank Dr. Xuehui Huang, Qi Feng, Danlin Fan and Congcong Zhou at the National Center of Plant Gene Research (Shanghai) for their excellent technique support. We thank Prof. Shuqun Cheng at the Eastern Hepatobiliary Surgery Hospital and Dr. Yumin Xu at the Ruijin Hospital for providing human liver tissues. We thank Ms. Mei Lu, Dr. Yalan Wu and all other members of the laboratory for their help.

## Author contribution

RH designed and supervised the study. YSL led the project; YSL, GHL, YFZ, KW and WDX performed the experiments. WDX, KW, SG, PW and MD performed mass spectrum and analyzed data. MHZ performed all the bioinformatic analyses. KW prepared the Samples for SILAC while WX and PX performed the proteomic analysis. JZ directed the synthesis of MDL8000 and instructed its use in animals. QD and YZ were responsible for all the animal work. RH, YSL, MHZ and GHL drafted the manuscript with inputs from all other authors. All authors read, and approved the final manuscript.

## Data and materials availability

All data needed to evaluate the conclusions in the paper are present in the paper and/or the Supplementary Materials. The generated data in this study have been deposited in the Gene Expression Omnibus (GEO, Accession No. GSE135860) and PRIDE (Accession No. PXD014908). To access the GEO data, using the secure: srqdeykspzmbhor. To access the PRIDE data, use this account, Password: jf8PgTab. Additional data related to this paper may be requested from the authors.

## Supplementary figures

**Figure S1:**
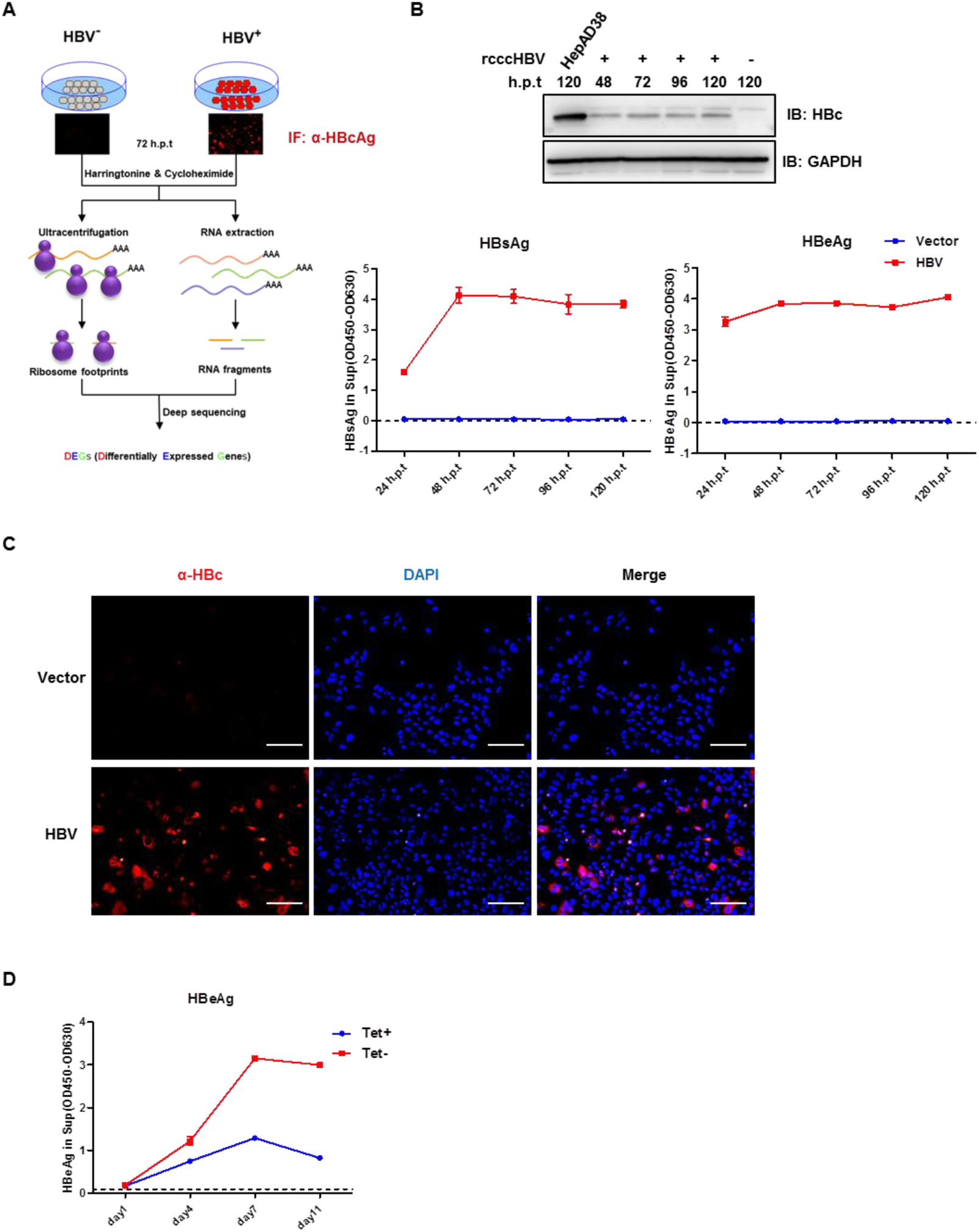
Replication kinetics of recombinant cccDNA system of hepatitis B virus. Huh7.5.1 cells were transfected with HBV recombinant cccDNA system (rcccHBV) or vector plus pCMV-Cre. (**A**) Experimental approaches for ribosome profiling of host genes upon transfection with the Cre-based rcccDNA system of HBV. (**B**) Cells were harvested at sequential time points as indicated, and lysates of HepAD38 cells which have been cultured for 120 hours after tetracycline withdraw was used as a positive control. h.p.t: hour post transfection. Culture medium in vector or HBV group was harvested and supernatants collected at sequential time points as indicated were subjected to ELISA (n=2). (**C**) Cells were subjected to immunofluorescence assay at 72 h.p.t. using anti-HBcAg (Dako, B0586, red) and nuclei stained by DAPI (blue). Scale bars, 100 μm. (**D**) The expression of HBeAg secreted from HepAD38 cells chromosomally integrated with the Tet-inducible HBV expression system at indicated time points after tetracycline withdraw was accessed by ELISA (n=2).

**Figure S2:**
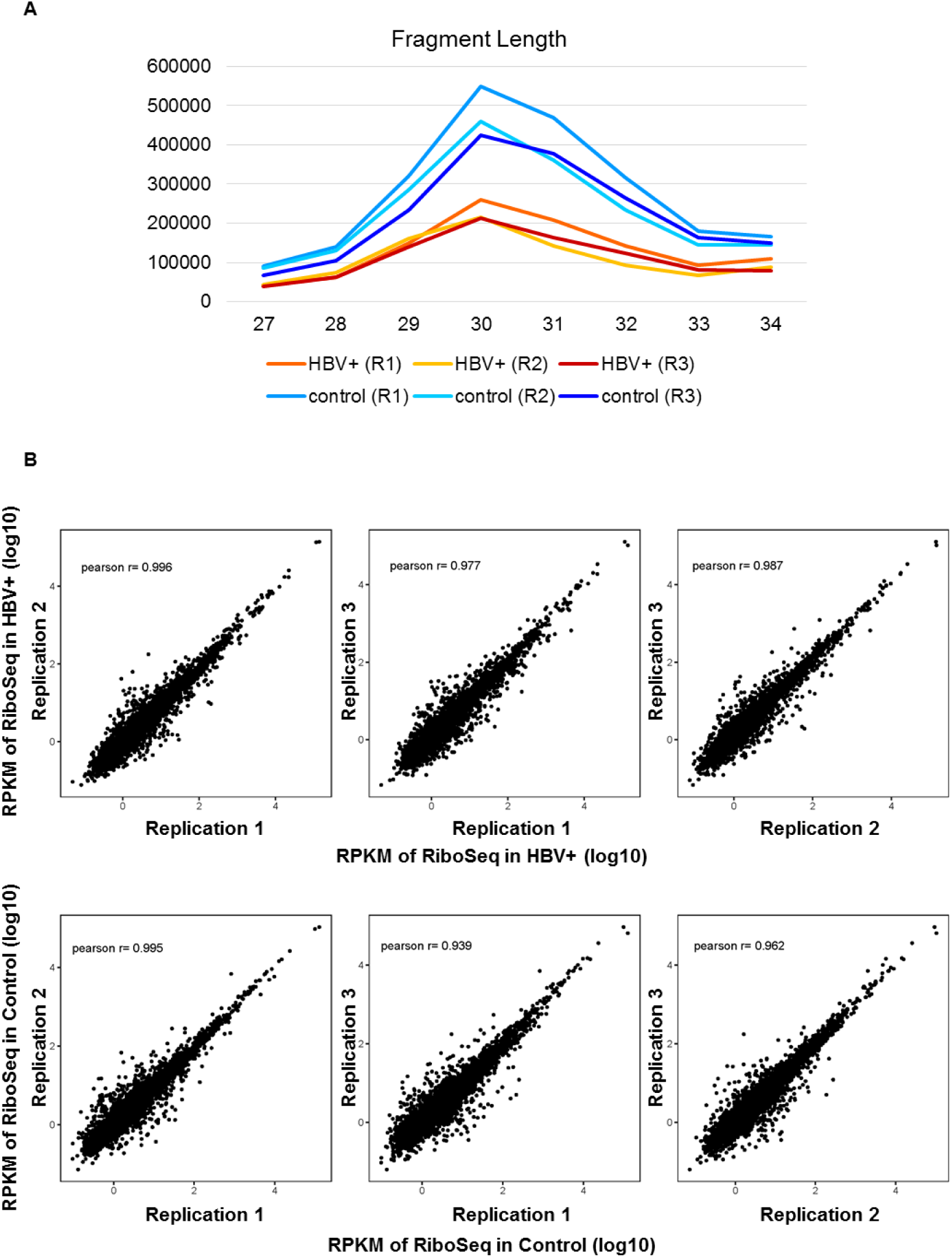
The length distribution of ribosome footprints and reproducibility of ribosome profiling experiments. (**A**) Control R1-3, replicate 1-3 in control group; HBV+ R1-3, HBV group replicates 1-3. X axis, the length distribution of ribosome footprints length in nucleotide (nt); Y axis, the reads numbers. (**B**) Plots show the correlations of RiboSeq RPKMs between three biological replicates in either HBV or non-HBV groups. Only mRNAs matched with > 5reads were counted.

**Figure S3:**
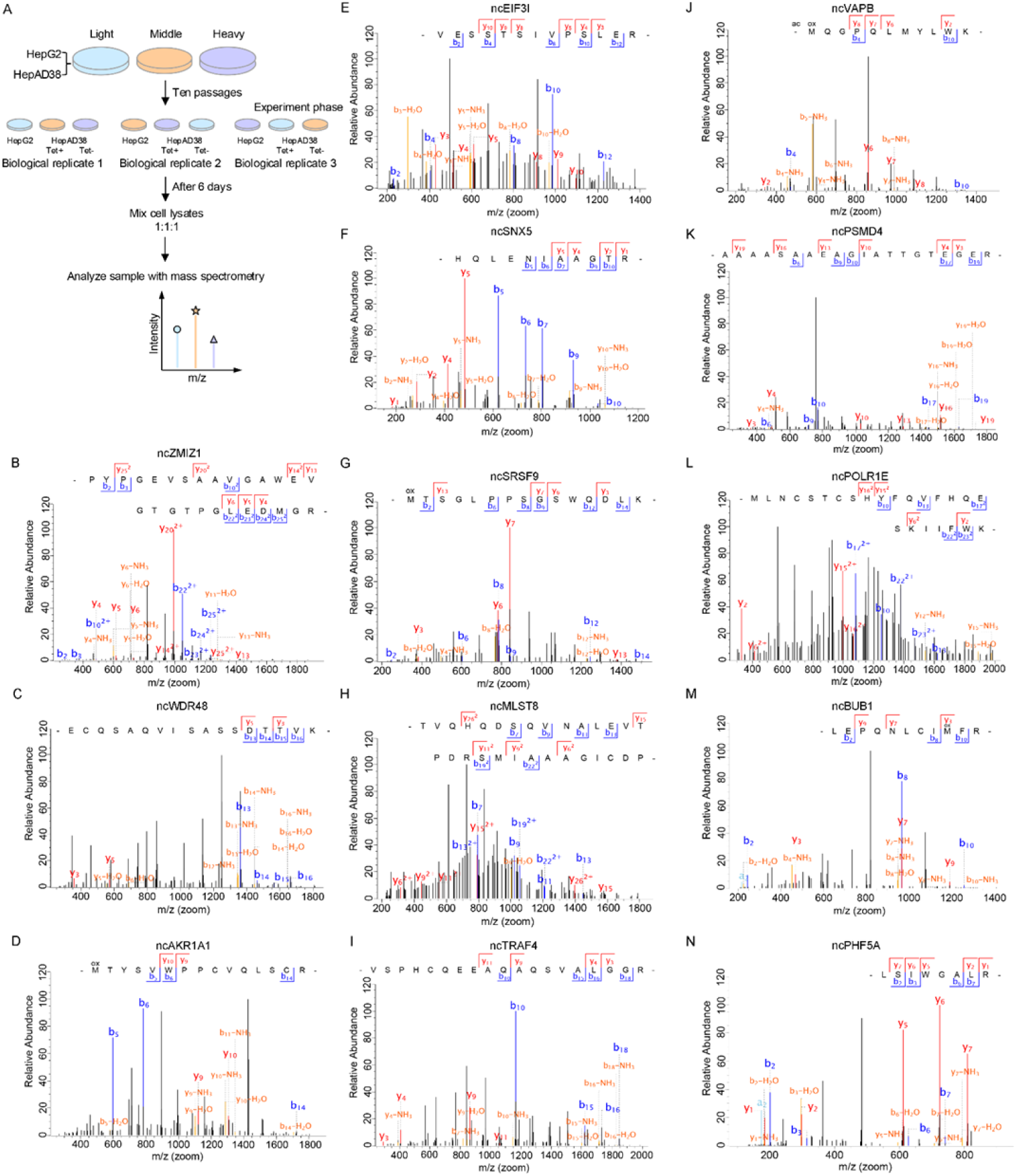
SILAC identified 13 peptides produced from non-canonical ORFs. (A) Three biological replicates were produced by labeling cells with different combination of light, middle and heavy stable isotopes, after experimental phase, cell lysates of each replicates were mixed by 1:1:1 and subjected to mass spectrometry (See online methods for more details). (B-N) MS spectra of identified peptide with sequence uniquely matched to the translated products of the non-canonical ORFs discovered in our ribosome profiling assays. These non-canonical ORFs were never annotated before and could have sequences and functions totally unrelated to any known proteins. They were nevertheless named after the closest annotated ORF in human genome. Matched y-ions and b-ions were shown in red and blue, respectively; the modified ions are shown in orange.

**Figure S4:**
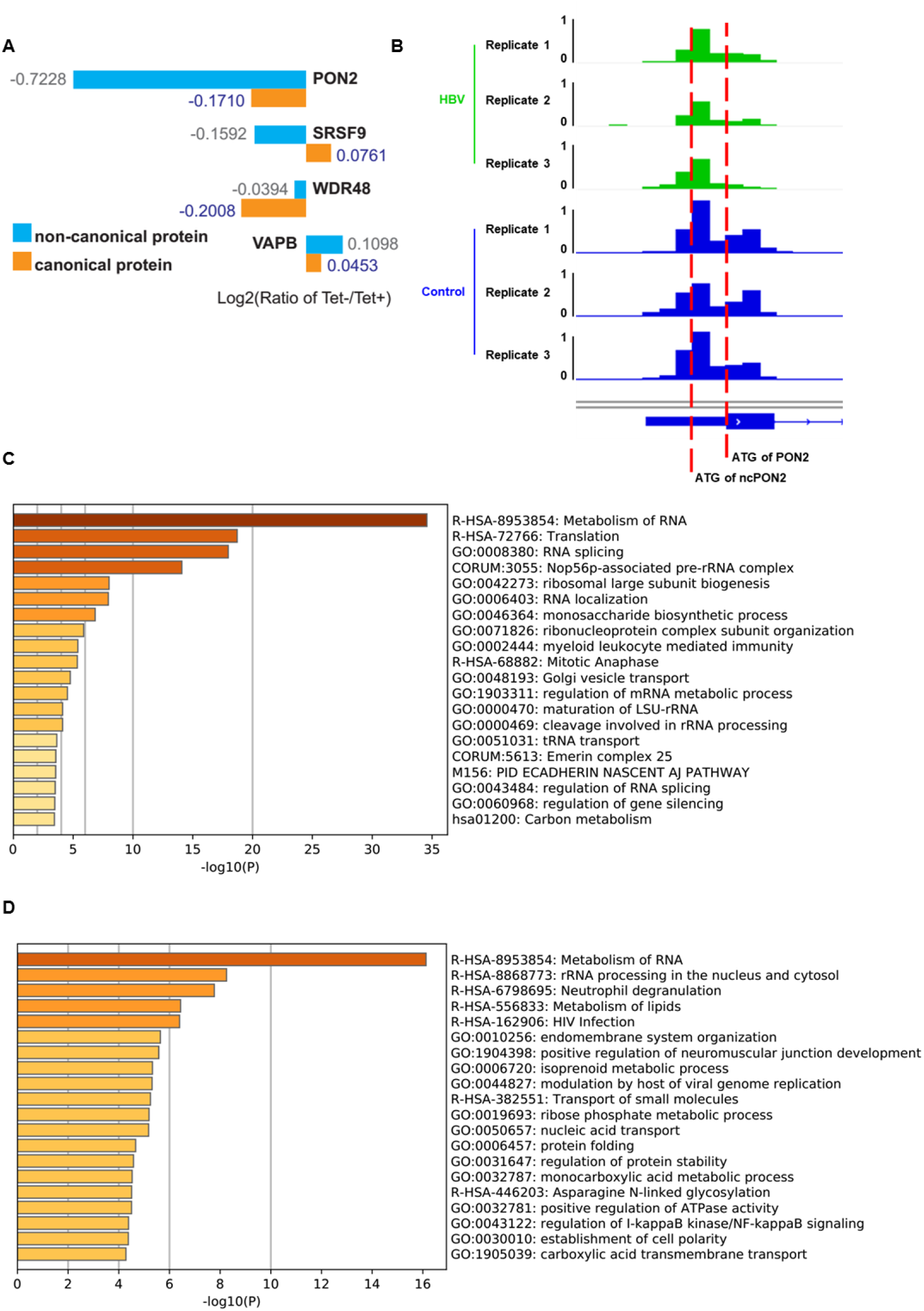
Analysis on ncPON2 and ncGRWD1. (**A**) The SILAC ratio of representative novel peptides and the corresponding canonical proteins in HBV+/HBV−, also see **Supplementary file S3 and S10**. (**B**)The IGV screen shot of RiboSeq pattern around the N-terminal region of *PON2* in all three biological replicates in either HBV or non-HBV groups. (**C** and **D**) To analyze the two HBV suppressive ncORFs, co-immunoprecipitation coupled with mass spectrometry analysis was performed with ncGRWD1-FLAG and ncPON2-FLAG, with pCDNA3.0-FLAG as a negative control. And GO and pathway enrichment analysis was performed using a web server Metascape (http://metascape.org/) with the identified interactors of ncGRWD1 (**C**) or ncPON2 (**D**) (also see **Supplementary file S4**).

**Figure S5:**
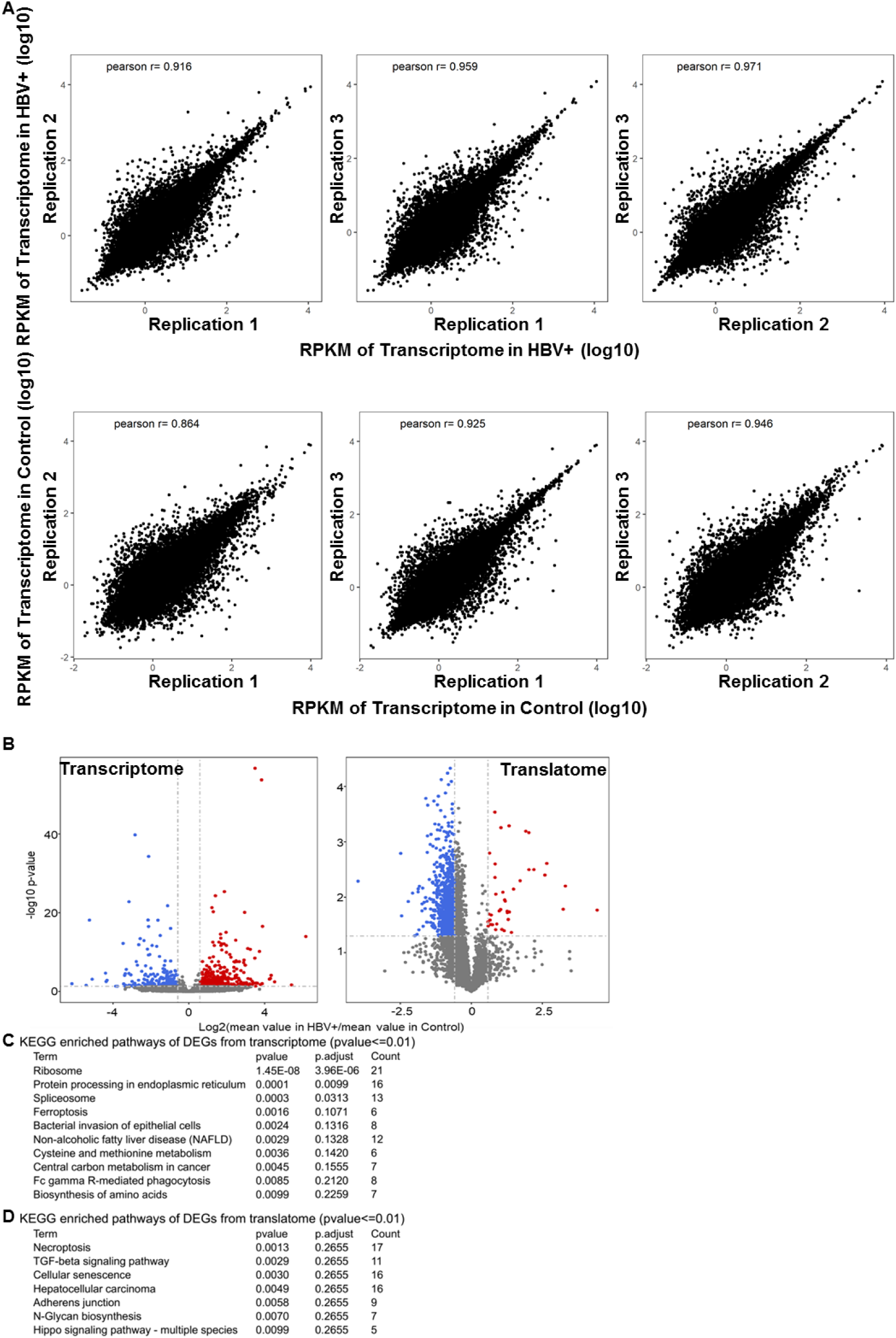
Reproducibility assay, volcano plot and KEGG pathway analysis of RNA-seq and ribosome profiling experiments. (**A**) Plots show the correlations of RNA-seq RPKMs between three biological replicates in either HBV or non-HBV groups. Only mRNA matched with > 64 reads were counted. (**B**) Differentially expressed genes (DEGs) in transcriptome and translatome were depicted as volcano plot, with *p*-value and fold change both shown; up-regulated and down-regulated genes were depicted as red and blue, respectively. (**C, D**) KEGG pathway enrichment analysis of genes differentially transcribed (**C**) or translated (**D**) in HBV-loaded cells versus control cells.

**Figure S6:**
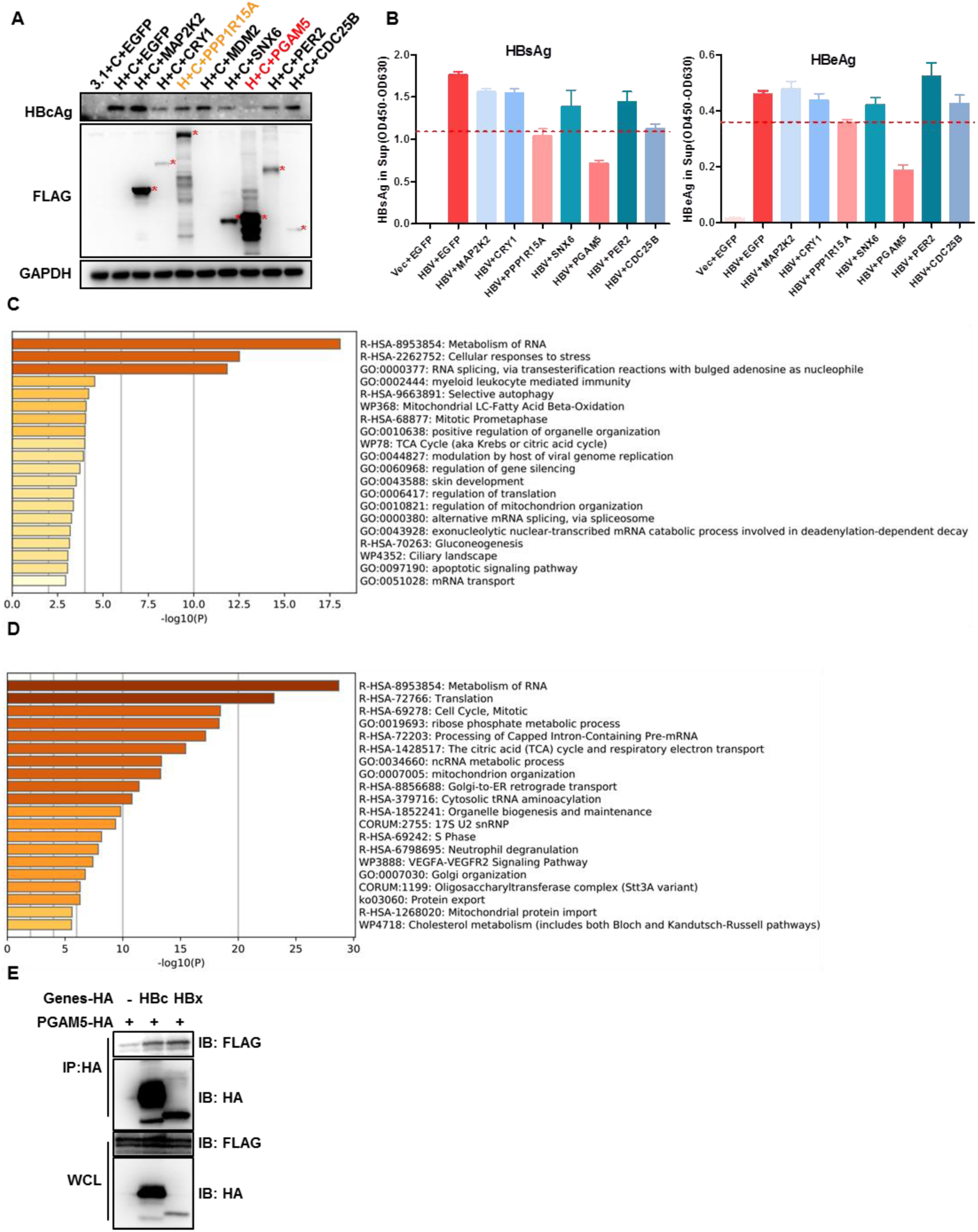
Screen and analysis on transcriptional and translational DEGs. **(A)** We tested the plasmids we have of the 35 transcriptional and translational DEGs as well as MDM2 in recombinant cccDNA system. Please note that we failed in detecting MDM2. (**C** and **D**) To further analysis the mechanism of the suppressive effect of PPP1R15A and PGAM5 on HBV, co-immunoprecipitation coupled with mass spectrometry analysis was performed with empty vector as a negative control to identify deemed interactions between host proteins and PPP1R15A or PGAM5, and GO and pathway enrichment analysis was performed using a web server Metascape (http://metascape.org/) with the identified interactors of PPP1R15A (**C**) or PGAM5 (**D**), also see **Supplementary file S9**. (**E**) We performed co-immunoprecipitation between PGAM5 and two proteins which were essential for virus replication, HBc and HBx.

**Figure S7:**
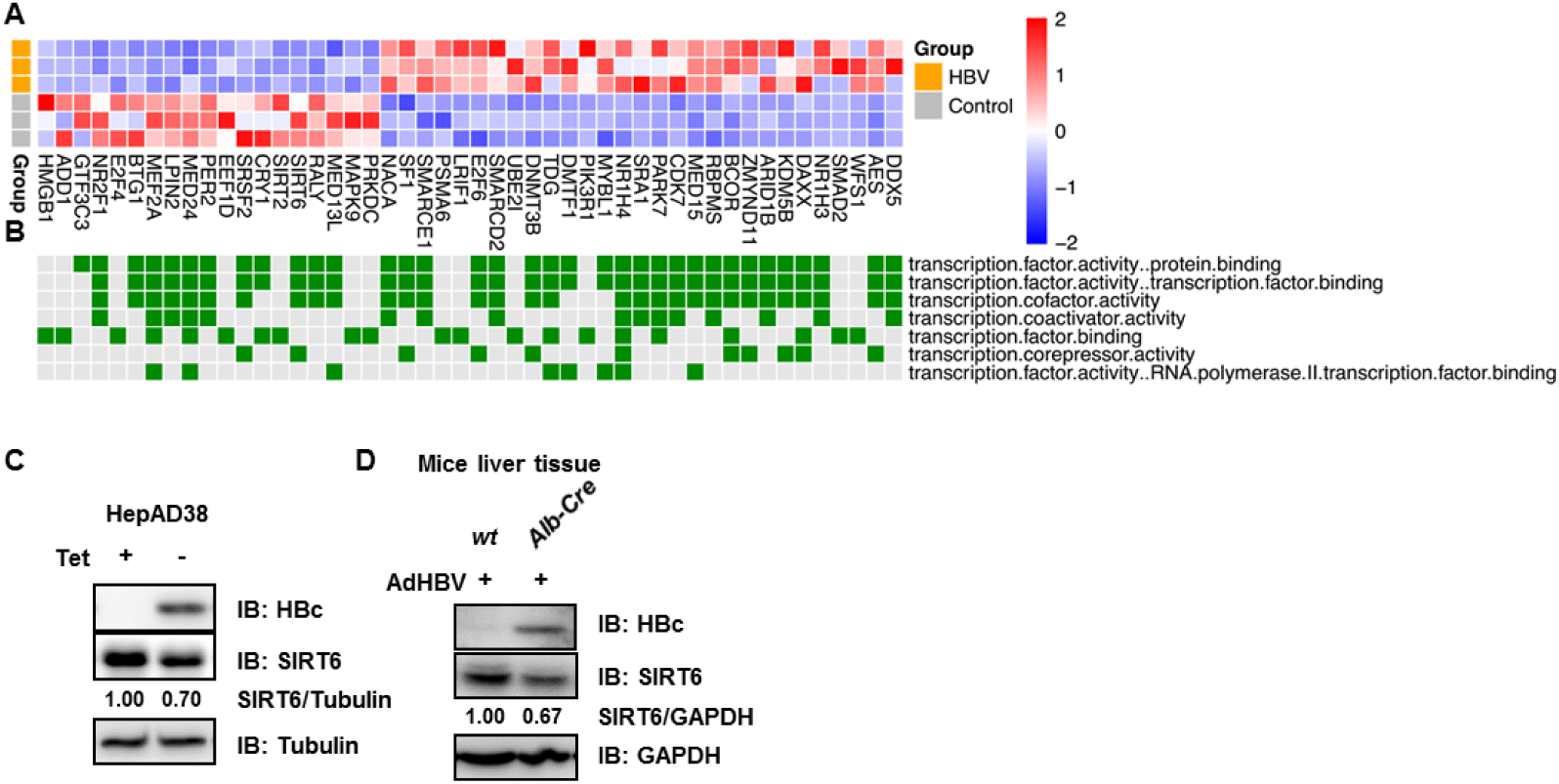
HBV down-regulates SIRT6 in HepAD38 cells and mouse model. (**A**) Heatmap of DEGs from RNA-seq in transcription molecular function. (**B**) The heatmap of DEGs participate in the indicated GO term. (**C**) HepAD38 cells with or without removal of tetracycline for 5 days were harvested, and lysates were subjected to IB analyses using indicated antibodies. (**D**) 14 days after recombinant adenovirus harboring HBV prcccDNA was injected into wild type or *Alb-Cre* transgenic mice, total proteins from the mouse liver tissues in each group were extracted and subjected to IB analyses using indicated antibodies.

**Figure S8:**
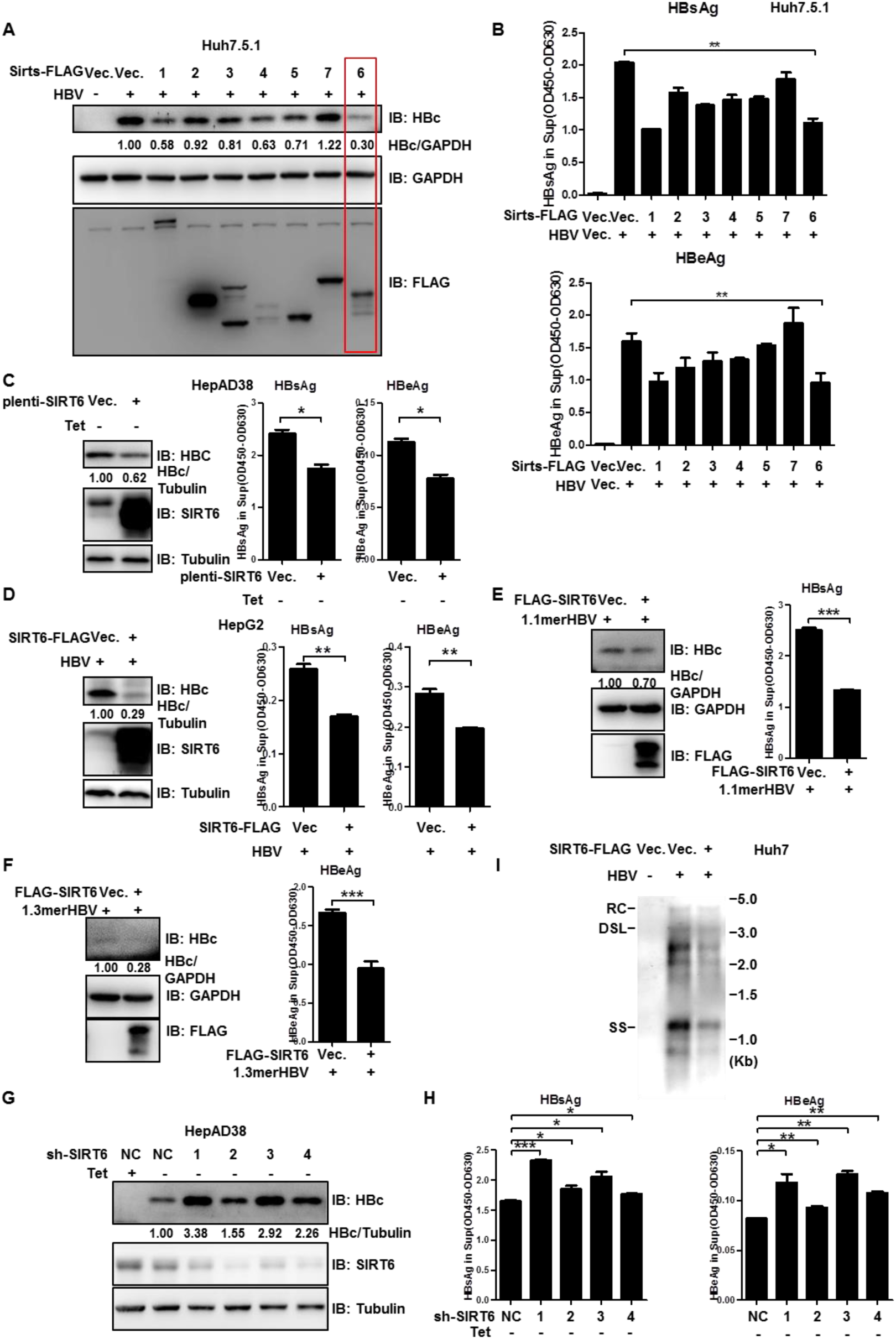
HBV down-regulates SIRT6 reciprocally in multiple HBV replication systems. (**A** and **B**)The cDNA of sirtuins-family were each co-transfected with HBV system into Huh7.5.1 cells and the protein levels of HBc, GAPDH and SIRTUINS were detected using indicated antibodies (A). The HBsAg and HBeAg level in supernatants were measured by ELISA. (B). (**C**) HepAD38 cells chromosomally integrated with the Tet-controlled HBV expression system were infected with lenti-virus vector expressing SIRT6, Cells were harvested and lysates subjected to IB using the indicated antibodies, while the supernatants from the cell culture were collected and subjected to ELISA using anti-HBsAg or anti-HBeAg. (**D**) HepG2 cells were co-transfected with HBV cccDNA system (H+C, prCCCDNA and pCMV-Cre) and pCDNA3.0-Sirt6-FLAG or pCDNA3.0 using Polyetherimide. Cells were harvested 72 h.p.t., with lysates collected and subjected to IB analyses using the indicated antibodies. The HBsAg and HBeAg in supernatants were determined by ELISA. (**E** and **F**) Experiments similar to those in D were performed with Huh7.5.1 cells transfected with 1.1mer- (E) or 1.3mer- (F) HBV linear genomes, For ELISA, n=3. (**G**) HepAD38 cells were transduced with lenti-virus vector (NC) or those encoding shRNAs that targeted endogenous SIRT6 (shSIRT6 1,2,3,4), After tetracyclin withdrawn for 4 days. HBcAg was measured using western blot, and (**H**) the level of HBsAg and HBeAg in supernatants was determined by ELISA. (**I**) The effect of SIRT6 on HBV genome replication was measured via southern blotting in Huh7 cells. RC, relaxed circular DNA; DSL, double strand linear DNA; SS, single strand DNA. *, p<0.05, **, p<0.01, ***, p<0.001.

**Figure S9:**
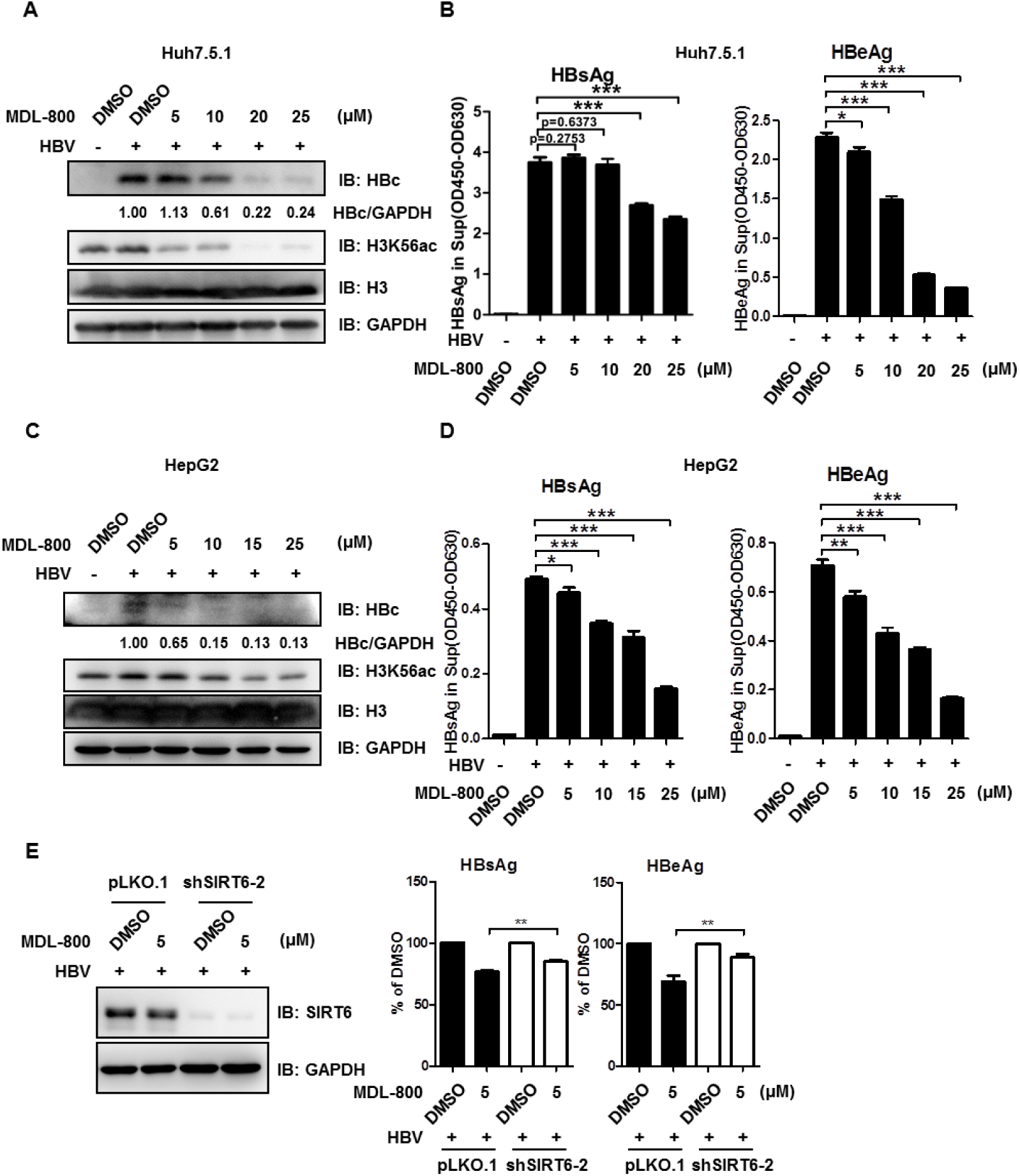
MDL-800 suppresses HBV gene expression in multiple cells. (**A-D**) Huh7.5.1 (A, B) or HepG2(C, D) cells were treated with increasing doses of MDL-800 after HBV transfection, the endogenous proteins were visualized with the indicated antibodies, while the levels of HBsAg or HBeAg in supernatants of cell cultures were determined with ELISA using anti-HBsAg or anti-HBeAg, n=3 for each group. (**E**) The effectiveness of MDL800 on HBV was tested in SIRT6 knock-down cell line or control cell line. For ELISA, n=3. *, p<0.05, **, p<0.01, ***, p<0.001.

**Figure S10:**
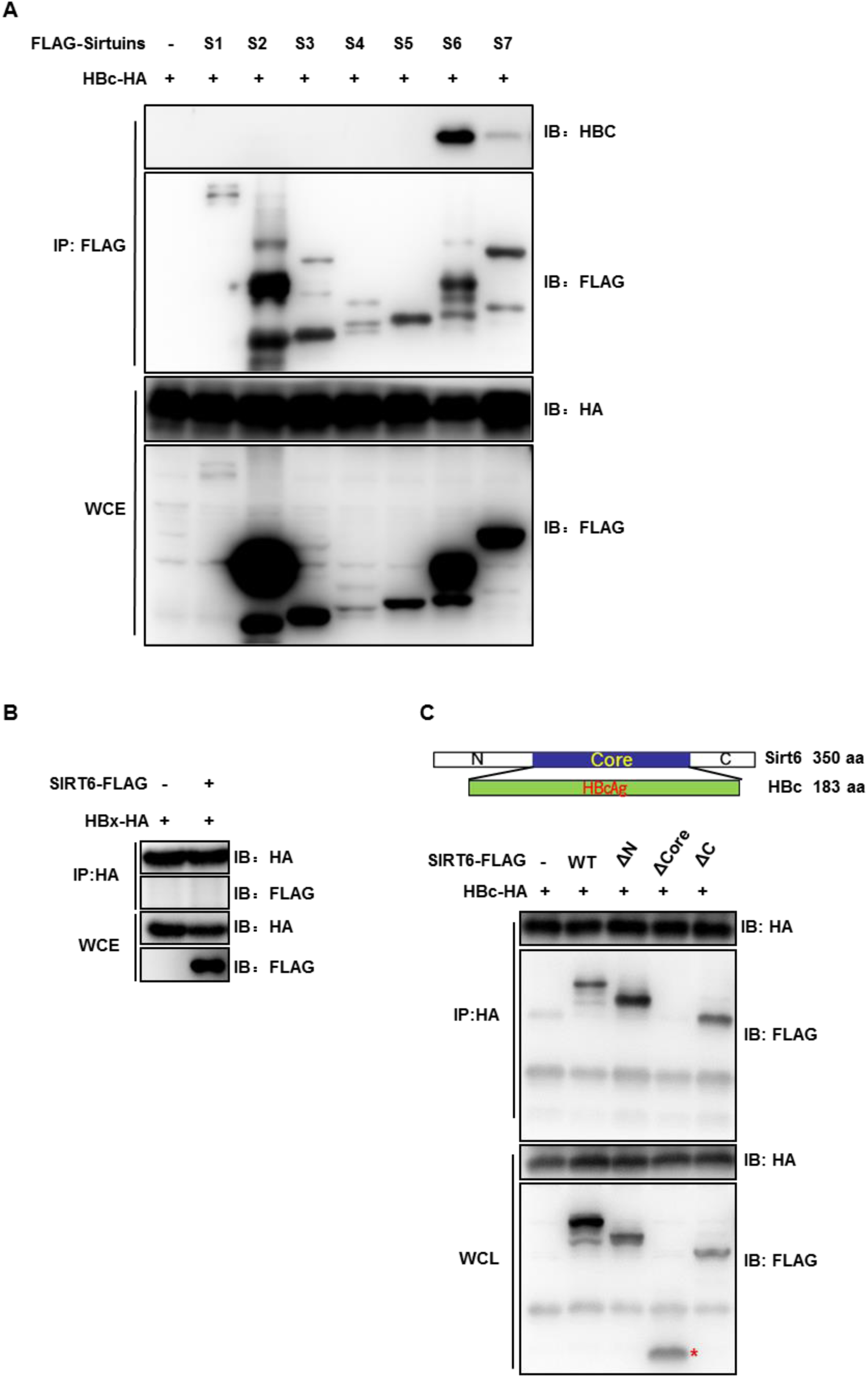
The interaction between SIRT6 and HBc. (**A**) HEK293T cells were transfected with HA-tagged HBc and FLAG-tagged Sirtuins family proteins, and co-immunoprecipitation was performed using anti-FLAG beads. (**B** and **C**) HEK293T cells were transfected with plasmids encoding HA-tagged HBx and FLAG-tagged SIRT6, with cells harvested at 48 h.p.t (hours post transfection), and lysates were subjected to Co-immunoprecipitation (Co-IP) followed by IB with indicated antibodies. (**B**) SIRT6 did not interact with HBx. (**C**) 293T cells were co-transfected with plasmids encoding HA tagged HBc and FLAG-tagged full-length SIRT6 or the indicated fragments, after 48 hours, Co-immunoprecipitation (Co-IP) was performed. **ΔN,** SIRT6 deleted the N terminal domain (1-48Aa); **ΔC**, SIRT6 deleted the C terminal domain (272-328Aa), **ΔCore**, SIRT6 with the catalytic core domain (49-271 Aa) deleted. WCE: whole cell extracts. WCL: whole cell lysates.

**Table S1:** Patient information in figure 3I and 3J. **Patient information in this study.** Five most widely used HBV test for HBsAg, HBsAb, HBeAg, HBeAb and HBcAb were determined at indicated hospital. HBV DNA copy number in some patient serum was also measured. Note that patient 5 and 6 were donors of liver transplantation and tested for HBV free.

| Patient Number | Clinical Diagnosis | Age | Gender | HBV DNA (copy/ml) | HBsAg (IU/ml) | Anti-HBs(mIU/ml) | HBeAg (S/CO) | Anti-HBe(S/CO) | Anti-HBc (S/CO) |
| --- | --- | --- | --- | --- | --- | --- | --- | --- | --- |
| 1 | Primary Liver Cancer Without HBV | 61 | Male | <50 | 0.01 | 0.01 | 0.35 | 0.08 | 0.5 |
| 2 | Liver Cirrhosis Without HBV | 58 | Male | NA | 0 | 108.94 | 0.28 | 0.34 | 7.38 |
| 3 | Liver Cirrhosis Without HBV | 48 | Male | NA | 0.03 | 0.69 | 0.26 | 0.03 | 9.23 |
| 4 | Liver Cirrhosis Without HBV | 71 | Male | NA | 0 | 3.54 | 0.33 | 1.42 | 7.92 |
| 5 | Donor of Liver Transplantation | NA | NA | 0 | 0 | NA | 0 | NA | NA |
| 6 | Donor of Liver Transplantation | NA | NA | 0 | 0 | NA | 0 | NA | NA |
| 7 | Primary Liver Cancer With HBV | 54 | Male | NA | 250 | 0.22 | 0.37 | 0.17 | 8.24 |
| 8 | Primary Liver Cancer With HBV | 66 | Male | <50 | 99.78 | 0.01 | 0.273 | 0.04 | 9.74 |
| 9 | Primary Liver Cancer With HBV | 51 | Male | <50 | 250 | 0.32 | 0.561 | 1.01 | 11.04 |
| 10 | Primary Liver Cancer With HBV | 53 | Male | 14500 | 250 | 0.06 | 0.294 | 0.01 | 11.79 |
| 11 | Primary Liver Cancer With HBV | 42 | Male | 8620 | 250 | <0.01 | 0.899 | 0.92 | 9.92 |
| 12 | Primary Liver Cancer With HBV | 46 | Male | 5990 | 250 | <0.01 | 0.335 | 0.01 | 12.02 |

Footnote: Patient 2-7 were diagnosed at Ruijin Hospital, others were diagnosed at Eastern Hepatobiliary Surgery Hospital. NA, Not Available; IU, International Unit; S/CO, Sample Optical Density / Cut-off Value.

**Data File S1. FPKM of ribosome profiling.**

**Data File S2. Ribocode analysis.**

**Data File S3. Novel peptides.**

**Data File S4. ncGRWD1 ncPON2 interactor.**

**Data File S5. FPKM of RNAseq.**

**Data File S6. DEGs list of RNAseq.**

**Data File S7. DEGs list of ribosome profiling.**

**Data File S8. Translation_efficiency.**

**Data File S9. PPP1R15A PGAM5 interactor.**

**Data File S10. proteome.**

**Data File S11. List of resources used in this study.**

